# PP4^Psy2^ regulates Rrm3 and Pif1 helicases to facilitate DSB resection across tandem retrotransposons

**DOI:** 10.1101/2025.02.19.639064

**Authors:** Adrián Campos, Lydia Pulido, Lydia Iglesias, Celia Delgado, Mar Sánchez, María Teresa Villoria, Andrés Clemente-Blanco

**Author notes:** Correspondence should be addressed to Andrés Clemente Blanco. These authors contributed equally to this work (Adrián Campos, Lydia Pulido).

## Abstract

Double-strand break repair is a tightly regulated process that relies on the coordinated activity of kinase and phosphatase complexes. Here we show that the PP4 phosphatase regulatory subunits, Psy2 and Psy4, act as positive and negative regulators of the holoenzyme during DSB repair. Cdk-dependent phosphorylation of Psy2 is crucial for DSB repair but is not required for DNA damage checkpoint inactivation. Lack of Psy2 leads to an asymmetric resection defect on one side of the DSB, a defect that is alleviated by expressing a phosphodeficient H2A variant or by depleting the checkpoint adaptor Rad9. This resection asymmetry is associated to the presence of tandem retrotransposons proximal to the DSB site, underlining the role of PP4^Psy2^ in stimulating resection across repetitive heterochromatic regions. PP4^Psy2^ support resection progression through paired transposable elements by reducing the phosphorylation state of Pif1 and Rrm3 helicases during the DSB response. These findings provide a comprehensive framework for the role of PP4^Psy2^ in resection-mediated DSB repair, highlighting its function in coordinating helicases activity to preserve genomic stability in response to DNA damage.

## Introduction

Cells are constantly exposed to different sources of DNA damage, which, if not properly addressed, can lead to the accumulation of genetic errors and genome instability. To preserve genomic integrity, cells have evolved highly conserved and sophisticated mechanisms to monitor the integrity of the DNA, detect lesions, and repair the damage, thereby preventing the propagation of mutations to the progeny. These processes, collectively known as the DNA damage response (DDR), activate the expression of DNA repair genes and initiate checkpoints that halt DNA replication and chromosomal segregation until the lesion has been repaired (Harper & Elledge, 2007). Among the most severe types of DNA damage are double-strand breaks (DSBs), which occur when both strands of the DNA double helix are simultaneously severed. DSBs can be repaired by either direct ligation of the broken ends through non-homologous end joining (NHEJ), or by homology-dependent mechanisms that rely on the searching of homologous sequences to regenerate the original DNA molecule in a process known as Homologous Recombination (HR) (Symington *et al*, 2014).

Homology-dependent DSB repair is always initiated by the nucleolytic degradation of the 5’-ended strand at both sides of the break to generate single-stranded DNA (ssDNA), a process known as resection (Cejka & Symington, 2021). Resection proceeds in two steps: an initial processing by the MRX complex (Mre11-Rad50-Xrs2) and Sae2, which creates short 3’ ssDNA overhangs (Gobbini *et al*, 2016), followed by extensive resection mediated by two redundant pathways involving the Sgs1/Dna2-Top3-Rmi1 complex and the Exo1 nuclease (Zhu *et al*, 2008). Extended resection generates long ssDNA tracts, which are coated by RPA to form a nucleofilament that engages the DNA damage checkpoint in order to coordinate DNA repair with cell cycle progression (Ciccia & Elledge, 2010). In *Saccharomyces cerevisiae*, the DNA damage checkpoint is primarily regulated by the phosphatidyl inositol kinase-like kinase (PIKK) Mec1, the yeast ortholog of ATR in mammals (Durocher & Jackson, 2001). Once activated, ssDNA-bound Mec1 transmits the checkpoint signal to other kinases such as Rad53 and Chk1. These kinases phosphorylate multiple downstream targets to orchestrate DDR processes, including a G2/M arrest that allows cells to accomplish the repair of the DNA lesion before cell cycle resumption (Waterman *et al*, 2020). To date, several distinct types of HR repair have been identified, each with unique characteristics, including synthesis-dependent strand annealing (SDSA), break-induced replication (BIR) and single-strand annealing (SSA) (Haber, 2016). SSA involves the alignment and annealing of complementary sequences flanking the break, typically facilitated by repeated sequences. After the DSB is processed to expose single-stranded DNA, the homologous regions anneal, and non-homologous ends are removed by nucleolytic cleavage. DNA polymerase and ligase then complete the repair. While SSA effectively restores DNA integrity, it is inherently mutagenic because it results in the loss of the intervening sequences between the homologous regions.

Protein phosphatase 4 (Ppp4/PP4/PPX) is a ubiquitous Ser/Thr phosphatase involved in multiple cellular functions independently of other protein phosphatases in the PPP family (Cohen *et al*, 2005; Ramos *et al*, 2019). PP4 was first identified in mammalian cells in a screening to isolate distinct forms of the PP1 phosphatase in various tissues (da Cruz e Silva *et al*, 1988). Structurally, PP4 shares 41% identity with PP1 and 65% with PP2A (Brewis & Cohen, 1992; da Cruz e Silva *et al*., 1988) and in *S. cerevisiae*, consists of a catalytic subunit, Pph3, and two regulatory elements, Psy2 and Psy4. Pph3 is not an essential protein, and contains high amino acid sequence similarity and functional resemblance to the PP2A catalytic subunits Pph21 and Pph22 (Ronne *et al*, 1991). The active phosphatase complex is formed by Pph3 and Psy2 (O’Neill *et al*, 2007), while Psy4 may provide substrate specificity. PP4 has been involved in multiple cellular processes, such as centrosome maturation (Helps *et al*, 1998; Sumiyoshi *et al*, 2002), spliceosomal assembly (Carnegie *et al*, 2003), cell signaling (Yeh *et al*, 2004), cell growth regulation (Bertram *et al*, 2000), chromatin remodeling (Zhang *et al*, 2005) and centromere pairing (Falk *et al*, 2010). The role of PP4 in the DDR was first evidenced through a high-throughput screen aimed to discover new proteins involved in DNA repair following UV irradiation (O’Neill *et al*, 2004). Further studies have shown that one of the main roles of PP4 in the DDR is to aid in checkpoint deactivation, enabling cell cycle reentry after repair. Specifically, PP4^Psy2^ has been involved in Rad53 dephosphorylation, facilitating cell cycle recovery after treatment with methyl methane sulfonate (MMS) (O’Neill *et al*., 2007). PP4 is also implicated in dephosphorylating γ-H2AX in diverse organisms, such as *Drosophila*, *S. cerevisiae* and humans, to promote checkpoint inactivation after DNA repair (Keogh *et al*, 2006; Merigliano *et al*, 2017; Nakada *et al*, 2008). Interestingly, the specificity of PP4 for dephosphorylating Rad53 and γ -H2AX is regulated by distinct binding subunits. PP4^Psy2^ mediates Rad53 dephosphorylation, as denoted by the hyperphosphorylation state of Rad53 in MMS-treated cells lacking Pph3 or Psy2. Conversely, PP4^Psy4^ does not contribute to Rad53 dephosphorylation following MMS exposure (O’Neill *et al*., 2007), but displays an increase in phosphorylated γ-H2AX levels (Keogh *et al*., 2006). Although these PP4 functions are primarily associated with checkpoint inactivation, in *S. cerevisiae*, PP4 also contributes to checkpoint activation by modulating the Mec1-Ddc2 complex (Hustedt *et al*, 2015). The Pph3-Psy2 complex physically interacts with Mec1-Ddc2 at replication fork collapse and DSBs sites, allowing coordinated regulation by kinase and phosphatase proteins over multiple DDR targets.

Beyond its role in DNA damage checkpoint regulation, PP4 is directly involved in the repair of DNA lesions. In mammals, the PP4C-PP4R2 complex (Pph3-Psy4 in yeast) participates in non-homologous end joining (NHEJ) repair of I-*SceI*-induced DSBs through its physical interaction with, and dephosphorylation of the chromatin condensation factor KAP1 (KRAB-associated protein 1) (Liu *et al*, 2012). In HR, PP4C-PP4R2 dephosphorylates the RPA2 subunit in response to replicative stress and DSBs, promoting RAD51 binding to resected DNA and facilitating recombination-based DNA repair pathways (Lee *et al*, 2010). Besides RPA2, PP4C is also essential for γ-H2AX dephosphorylation, further stimulating HR (Chowdhury *et al*, 2008; Nakada *et al*., 2008). Supporting these findings, PP4-mediated Rad53 dephosphorylation in *S. cerevisiae* enhances DNA resection by counteracting the inhibitory effect of Rad9 on the Dna2-Sgs1 complex (Villoria *et al*, 2019).

Notably, PP4 regulation is influenced by post-transcriptional modification of its regulatory subunits. Psy4 undergoes CDK-dependent phosphorylation, which controls its subcellular localization during the cell cycle. This phosphorylation occurs specifically at Thr320 and Thr347, leading to a rapid cytoplasmic re-localization of Psy4 during S-phase (Faustova *et al*, 2022; Kosugi *et al*, 2009). Additionally, phosphoproteomics studies have identified several phosphorylation sites on Psy2 (Lanz *et al*, 2021), although the exact kinases responsible for these modifications and its function remains unknown.

Here we show that Pph3 regulatory elements Psy2 and Psy4 act as positive and negative regulators of PP4 in response to a DSB. Cdk-dependent regulation of Psy2 is essential for efficient SSA but not for checkpoint deactivation. Psy2 is crucial for sustaining asymmetric resection at the DSB vicinity by facilitating resection across retrotransposons sequences. This role in stimulating end resection throughout repetitive genomic regions is achieved by modulating Pif1 and Rrm3 helicases phosphorylation levels during the damage response. Overall, these results establish a new function for PP4^Psy2^ in DNA repair and highlight its critical role in regulating resection and helicase activity to ensure efficient repair of DSBs, particularly in repetitive genomic regions.

## Results

### PP4 regulatory elements Psy2 and Psy4 have opposite functions in DNA repair by SSA

Deletion of *PPH3*, which encodes the catalytic subunit of the phosphatase complex PP4, has been shown to delay the repair of an HO-inducible DSB through Single Strand Annealing (SSA). Biochemically, PP4 modulates the DNA damage response by buffering Rad53 phosphorylation levels, thereby reducing histone H2A phosphorylation and the binding of the adaptor protein Rad9 to chromatin surrounding the HO break site. This PP4-dependent attenuation of the DNA damage checkpoint is essential for efficient resection by the Sgs1/Dna2 complex during the early stages of the repair (Villoria *et al*., 2019). To dissect the specific roles of the individual PP4 regulatory subunits in controlling Pph3 in response to a DSB, we generated single *pph3Δ*, *psy2Δ* and *psy4Δ* mutants, and assessed SSA efficiency in the *YMV80* background (Fig 1A). This background contains an HO target site integrated into the *LEU2* gene on chromosome III (*LEU2::cs*) (Fig 1A). The HO recognition site at the endogenous *MAT*a locus was deleted to prevent the formation of a DSB at this site upon HO expression (Fig 1A). The endogenous HO promoter was replaced with the galactose-inducible *GAL1* promoter, restricting DSB induction at the *LEU2::cs* site only when galactose is added to the medium. As donor for repair, a 5’ fragment of the *LEU2* gene (referred to as *U2*), was inserted left to the HO site (Fig 1A). Upon galactose addition, the resulting DSB can be repaired either by Rad51-independent Single Strand Annealing (SSA) or Rad51-dependent Break Induced Replication (BIR) pathways. Given the known role of PP4 in promoting DNA end-resection, and the critical contribution of resection to SSA, we employed *RAD51*-disrupted strains to bias DNA repair toward SSA (Fig 1A).

**Figure 1.**
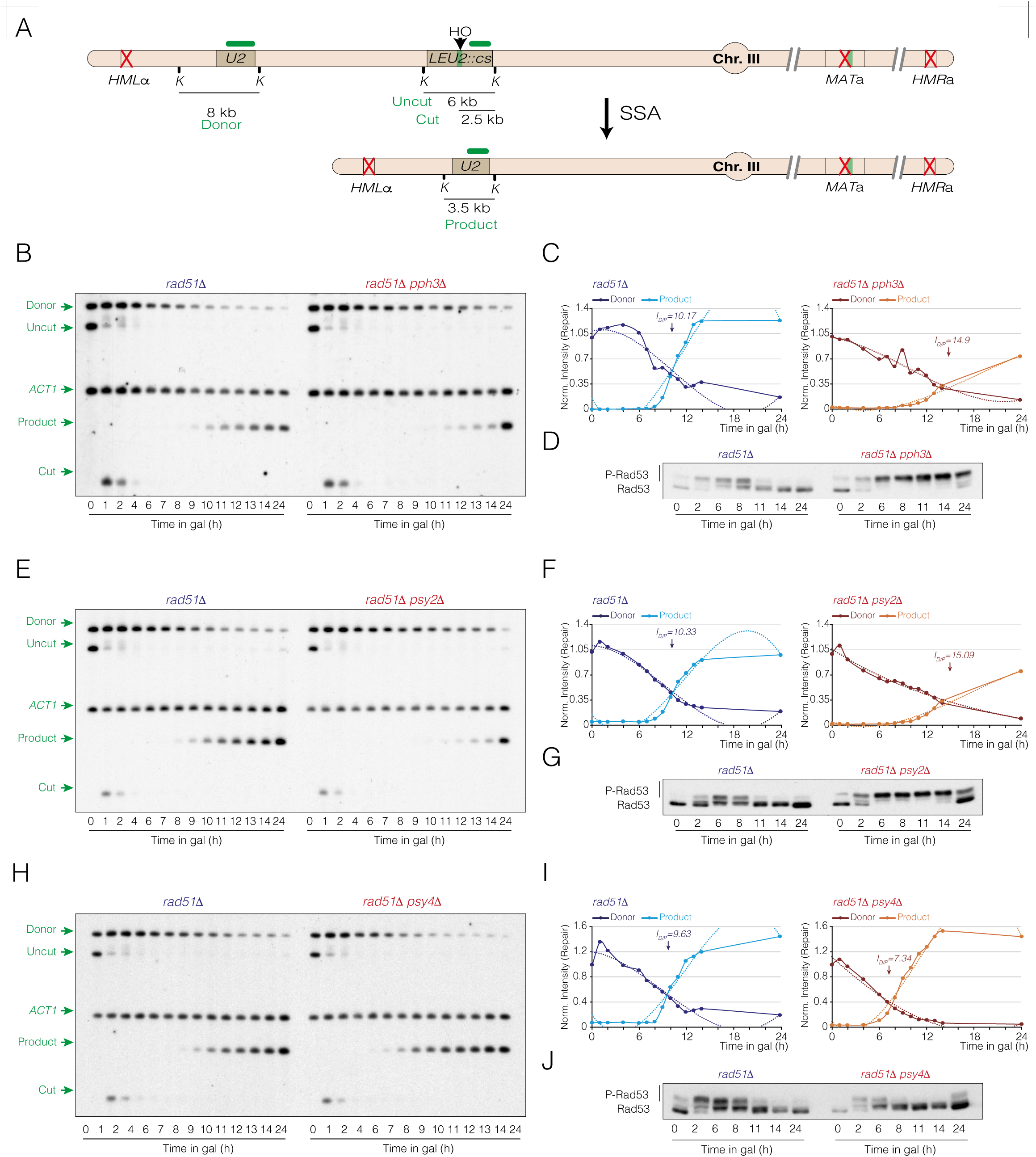
The PP4 regulatory subunits Psy2 and Psy4 exhibit opposing roles in DNA repair by SSA. **A.** Schematic representation of the relevant genomic structures of the *YMV80* strain. The *MAT*a, *HML*a and *HML*α *loci* on chromosome III have been deleted to prevent inter-chromosomal SDSA recombination at the *MAT locus*. The positions of the HO cleavage site at the *LEU2 locus* (*LEU2::cs*) and the donor sequence (*U2*) are illustrated. The sizes of the donor, uncut, cut, product bands, as well as the distances used to assess DNA repair dynamics, are depicted. The localization of the probes (horizontal green lines) and the restriction endonuclease cleavage sites (K: *Kpn*I) used for Southern blot analysis are also shown. **B.** Southern blot of *rad51Δ* and *rad51Δ pph3Δ* cells carrying the DNA repair system depicted in A. Cells were grown overnight in YP-Raffinose before inducing the HO nuclease with galactose. Samples were collected at various time points, genomic DNA was extracted, digested with *Kpn*I, and analyzed by Southern blot using the probes indicated in A. The *ACT1* gene sequence was used as loading control. **C.** Graphs derived from the data obtained in B, representing the quantification of the averaged donor and product band signals from two biological replicates, normalized against their respective *ACT1* and uncut T0 band signals. The intersection point of the donor and product trend lines (I_D/P_ value) represents the mean from two independent experiments. The statistical significance (*P*-value) of differences between both strains assessed by a two-tailed unpaired Student’s *t-*test is 0.0042 (**). **D.** Samples from the experiment shown in B were collected at the indicated time points, and proteins were extracted using TCA precipitation. Western blot analysis was performed to detect distinctives Rad53 phospho-isoforms along the HO induction. **E.** Southern blot analysis of *rad51Δ* and *rad51Δ psy2Δ* cells conducted under the same experimental conditions as in B. **F.** Graphs obtained from the data in E, showing the quantification of averaged donor and product band signals from two independent experiments, normalized against their respective *ACT1* and uncut T0 band signals. The intersection point of the donor and product trend lines (I_D/P_ value) represents the mean from two independent experiments. The statistical significance (*P*-value) of differences between both strains assessed by a two-tailed unpaired Student’s *t-*test is 0.0378 (*). **G.** Samples from the experiment shown in E were collected at the indicated time points, proteins were TCA extracted and subjected to western blotting analysis to evaluate Rad53 phosphorylation levels along the experiment. **H.** Southern blot analysis of *rad51Δ* and *rad51Δ psy4Δ* cells performed under the same experimental conditions as in B. **I.** Graphs obtained from the analysis of the data in H, representing the quantification of averaged donor and product band signals from two biological replicates after normalizing against their respective *ACT1* and uncut T0 band signals. The intersection point of the donor and product trend lines (I_D/P_ value) represents the mean from two independent experiments. The statistical significance (*P*-value) of differences between both strains assessed by a two-tailed unpaired Student’s *t-*test is 0.604 (ns). **J.** Samples from the experiment in H were collected at the indicated time points, proteins were TCA extracted and subjected to western blotting analysis to detect Rad53 phospho-isoforms.

To evaluate the impact of Pph3 on SSA efficiency, we induced HO expression in exponentially growing cells of *rad51Δ* and *rad51Δ pph3Δ* mutants by adding galactose to the medium. Samples were collected at various time points and analyzed by Southern blot using a probe targeting the *U2* sequence (Fig 1A). In both strains, the 6 kb uncut band signal decreased dramatically, accompanied by the appearance of the 2.5 kb cut product within one hour from the HO induction, indicating a rapid and synchronous HO break induction in both strains (Fig 1B). No differences were observed in the disappearance kinetics of the cut fragment between both strains, denoting comparable initial resection rates. However, a notable delay was observed in the *rad51Δ pph3Δ* strain in the disappearance of the donor band and the accumulation of the 3.5 kb repair product compared to the single *rad51Δ* mutant (Fig 1B). These results confirm that absence of Pph3 impairs SSA efficiency under these experimental conditions.

In the context of SSA, resection through the *U2* donor sequence is a prerequisite for the formation of the repair product. Consequently, the disappearance of the *U2* donor band always precede the accumulation of the repair product signal. To accurately measure the dynamics of DNA end resection, we calculated the time at which the donor and product trend lines intersect, a time-point that we refer as to I_D/P_ value (Fig 1C). This value represents the average time required for resection to reach the *U2* sequence and repair the HO break, providing a reliable metric for assessing resection and repair dynamics in a given cell population. We measured an I_D/P_ value of 10.17 h in the control *rad51Δ* strain, which increased to 14.9 h in the absence of Pph3 (Fig 1C). Consistent with previous studies (O’Neill *et al*., 2007; Villoria *et al*., 2019), we detected prolonged phosphorylation of Rad53 in *rad51Δ pph3Δ* cells compared to the *rad51Δ* control strain (Fig 1D). Similarly, SSA dynamics in *psy2Δ* mirrored those in *pph3Δ* cells (Fig 1E), with a similar I_D/P_ value of 15.09 h (Fig 1F), and sustained levels of Rad53 phosphorylation throughout the experiment (Fig 1G). These findings suggest that Psy2 acts as a positive regulator of Pph3 during SSA repair. Accordingly, a double *psy2Δ pph3Δ* mutant retrieved similar DNA repair kinetics than a single *psy2Δ* mutant (Fig EV1A), with only a slight increase in the I_D/P_ value from 14.47 h to 16.18 h (Fig EV1B). Supporting this result, the dynamics of Rad53 phosphorylation were similar between the two strains (Fig EV1C). Conversely, *psy4Δ* cells showed a slightly faster disappearance of the donor band and earlier accumulation of the repair product (Fig 1H). This corresponded to a reduced I_D/P_ value of 7.34 h compared to 9.63 h in the *rad51Δ* control strain (Fig 1I) and a pronounced attenuation of Rad53 phosphorylation levels (Fig 1J). These results indicate that Psy4 functions as a negative regulator of the PP4 holoenzyme during SSA-mediated DNA repair.

### Cdk phosphorylation of Psy2 is required for SSA repair but not for PP4-dependent checkpoint inactivation

We previously stablished that Psy2 functions as a positive regulator of Pph3 in SSA. To investigate whether this regulatory role involves post-transcriptional modifications of the protein, we looked for changes in its phosphorylation profile during the DNA damage response. Induction of an HO break led to a transient accumulation of high molecular weight isoforms of Psy2 in Phos-tag western blots experiments during the early stages of the damage response, indicative of DNA damage-dependent phosphorylation events on the protein (Fig EV1D). Expression of a *cdc28-as1* allele reduced these higher molecular weight isoforms following the addition of 1-NM-PP1 but not in mock DMSO (Fig EV1E). These results demonstrate that Psy2 undergoes Cdk-dependent phosphorylation in response to DNA damage. The amino acid sequence analysis of Psy2 revealed four Cdk consensus sites located at the N-terminal domain: T6, S10 T12 and S68 (Fig EV1F). Expression of a phospho-deficient *psy2-S/T-A* version lacking these four Cdk consensus sites in *rad51Δ* cells impaired SSA, as denoted by the delay in the disappearance of the donor band and in the accumulation of the repair product compared to the *rad51Δ* control strain (Fig EV1G). Supporting these findings, the I_D/P_ value slightly increased from 10.69 h in control *rad51Δ* cells to 12.25 h in *rad51Δ psy2-S/T-A* cells (Fig EV1H). Interestingly, the pattern of Rad53 phosphorylation during the HO response was similar between both strains (Fig EV1I), suggesting that PP4*^psy2-S/T-A^*affects SSA in a Rad53-independent manner.

### Genome-wide sequencing analysis of SSA reveals a Psy2-dependent resection symmetry

The analysis of the reference genome provided by the *Saccharomyces* Genome Database (SGD) revealed a unique retrotransposable element (*YCLWTy2-1*) located to the left of the *LEU2* locus, and an expected inter-*U2* distance of 22.5 kb. To accurately measure resection dynamics in the *YMV80* strain, we performed long-read sequencing and assembled the region spanning the *LEU2::cs* and *U2* loci. This assembly revealed an additional Ty element located left to the previously described *YCLWTy2-1* retrotransposon (Fig 2A). FASTA alignment of this Ty element against the *S. cerevisiae* genome showed the highest similarity to the Ty *YERCTy1-1* at chromosome V, indicating a retrotrasposition event in the *YMV80* background (Fig 2A). We constructed a new reference genome (hereafter referred to as *YMV80_v2*) incorporating the tandem *YERCTy1-1/YCLWTy2-1* module, which was used for subsequent sequencing read alignments throughout this study.

**Figure 2.**
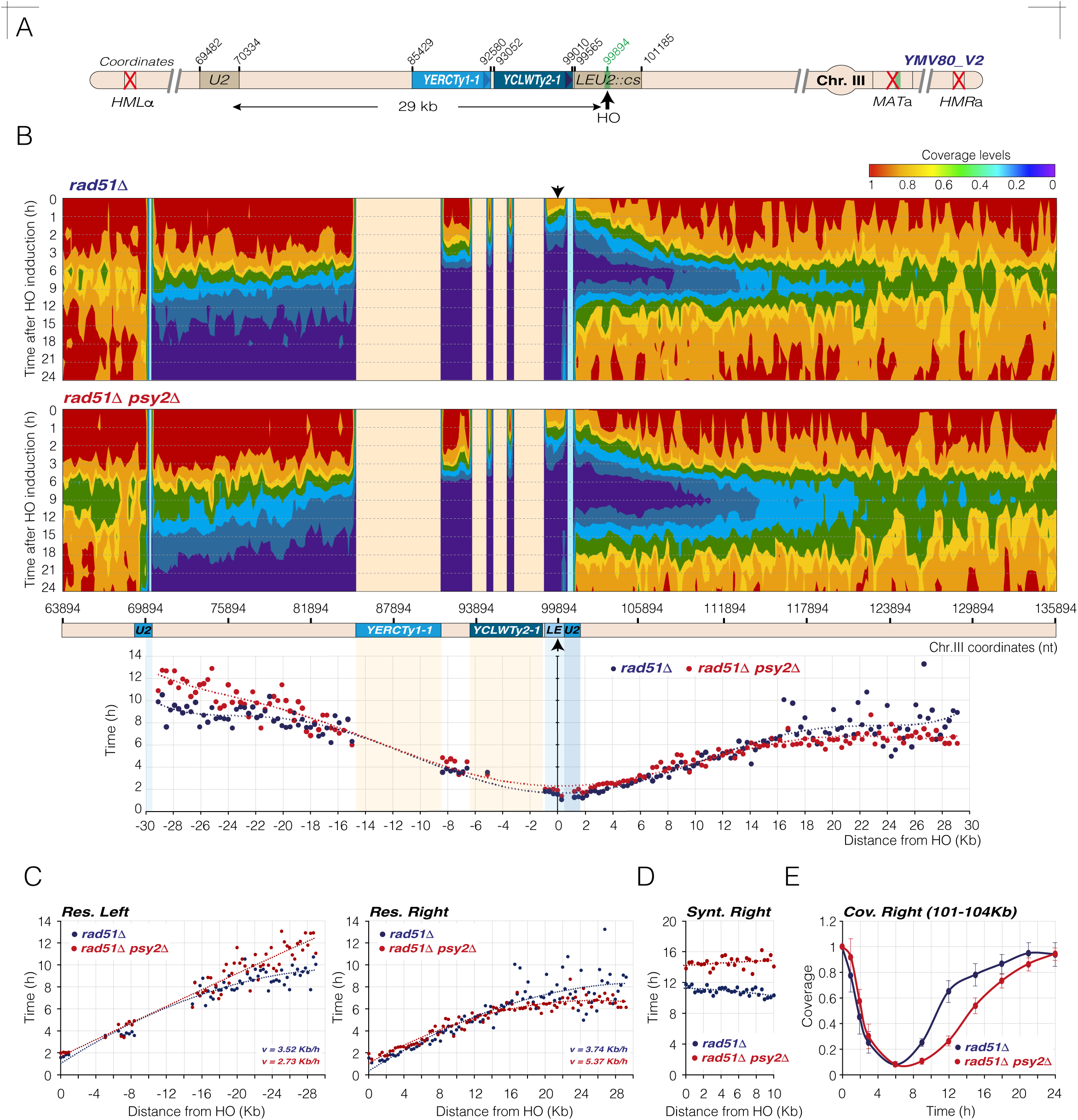
Genome-wide sequencing analysis of SSA reveals a Psy2-dependent resection symmetry. **A.** Schematic representation illustrating the distinctive genomic features of the *YMV80_v2* strain, as determined by long-read sequencing assembly. The coordinates for each feature and the distance between the two *U2* homologous sequences are shown. **B.** Colormap charts displaying coverage levels in *rad51Δ* and *rad51Δ psy2Δ* strains. Coverage levels, normalized against the T0 sample, are represented by red, green and blue gradients. The *x-*axis corresponds to chromosome III coordinates, while the *y-*axis represents time after HO induction. Black arrows mark the position of the HO cleavage site. Data shown are the mean of two biological experiments. Note that repetitive Ty elements (red areas) and *U2* sequences (blue areas) were excluded from the analysis. Bellow, a graph shows time (h) *versus* distance (kb) across a +/-30 kb region from the HO site. This was assessed by analyzing the trend line equations for 300 nt average coverage fractions as they dropped from 1 to 0.5 (see Materials and Methods for details). **C.** Analysis of the resection dynamics in *rad51Δ* and *rad51Δ psy2Δ* cells. The left and right sides of the HO site from the graph depicted in B were split and analyzed separately. Trend line equations were derived from the time (*y*-values) required for resection to cover distinctive distances (*x*-values) up to 30 kb from the HO break. Resection rates (*v*) were calculated by measuring the time (*Δt*) needed for resection to accomplish this distance (*Δs*) at both sides of the break (see Materials and Methods for details). **D.** Determination of DNA re-synthesis rate in *rad51Δ* and *rad51Δ psy2Δ* mutants. Synthesis rates were calculated by averaging coverage levels of 300 nt fractions spanning from 0 kb to 10 kb to the right of the HO site during the 9 h – 24 h interval. The trend line equations for each fraction were used to determine the time (h) required for coverage values to recover to 0.5 (see Materials and Methods for details). **E.** Analysis of coverage levels on the right side of the HO site in *rad51Δ* and *rad51Δ psy2Δ* strains during HO nuclease induction. The coverage of a right HO region between coordinates 101,000 and 104,000 on chromosome III was averaged and plotted over time.

To thoroughly characterize the role of PP4^Psy2^ in DNA resection and repair by SSA, we adapted the GWS pipeline previously applied for analyzing inter-chromosomal SDSA repair in the *PMV* background (Campos *et al*, 2023; Ramos *et al*, 2022) to the *YMV80_v2*. In this approach, only double-stranded DNA fragments are incorporated into sequencing libraries, allowing us to infer the length, directionality, and dynamics of resection by analyzing changes in coverage near the HO cut site following its induction. The HO nuclease was induced with galactose in exponentially growing *rad51Δ* and *rad51Δ psy2Δ* cells, and samples were collected at various time points for genome-wide sequencing. Sequencing reads were aligned to the *YMV80_v2* reference genome, and coverage levels were normalized to undamaged T0 samples. This approach enabled the generation of two-dimensional (2D) (Fig EV2A), three-dimensional (3D) (Fig EV2B) and heatmaps (Fig 2B) coverage profiles, providing detailed insights into the progression of the HO-induced break over time.

Analysis of the coverage profile in the *rad51Δ* mutant revealed a gradual and symmetrical loss of coverage on both sides of the break during the 1-9 h interval following HO expression (Fig 2B, top panel, Fig EV2A, left graphs and Fig EV2B, left graph). By 9 h post-induction resection had extended to the *U2* donor sequence. Note that the *U2* (blue areas) and *YCLWTy2-1/YERCTy1-1* (orange areas) sequences were excluded from the analysis, as their repetitive nature impairs the correct assessment of reads aligning at these genomic regions. Once resection reached the *U2* donor sequence, cells rapidly recovered coverage levels on the right side of the HO break, due to the pairing of the two homologues *U2* sequences and subsequent polymerization of the resected tracts by damage-specific DNA polymerases (Fig 2B, top panel, Fig EV2A, left graphs and Fig EV2B, left graph). As expected, SSA repair left a distinct gap with minimal coverage between the two *U2* sequences, reflecting the nature of the DNA repair mechanism.

Consistent with the results from Southern blot experiments (Fig 1E), the *rad51Δ psy2Δ* mutant exhibited defects in DNA end resection on the left side of the HO break, as evidenced from the slower reduction in coverage levels as resection progresses toward the *U2* donor sequence. Interestingly, resection on the right side of the HO break was not delayed, indicating that the resection defect in *rad51Δ psy2Δ* cells is asymmetric, specifically affecting the left side of the cleavage site (Fig 2B, bottom panel). As anticipated, the reduced resection rate on the left side of the HO break in *rad51Δ psy2Δ* cells resulted in a significant delay in the recovery of coverage levels on the right side of the break (Fig 2B, bottom panel, Fig EV2A, right graphs and Fig EV2B, right graph).

To quantitatively measure resection rates on both sides of the HO break in *rad51Δ* and *rad51Δ psy2Δ* strains, we averaged the sequencing coverage across 300 nt fractions spanning a +/-30 kb region around the HO site for each time-point analyzed. Using the trend line equations generated for these fractions, we calculated the time required for coverage to drop from 1 to 0.5. This metric reflects the averaged distance that resection has reached in the cell population at a given time point. It is important to remark that in SSA, the DNA region between the homologous *U2* sequences is not subjected to DNA re-synthesis. Therefore, the decrease in coverage during HO induction directly correlates with resection progression. Accordingly, we used a 0-15 h time interval to generate the trendlines equations when dissecting the left side of the HO break. However, since at the right side of the HO site coverage levels are influenced by both resection and DNA re-polymerization, we limited the analysis to the 0-6 h interval. This avoids the potential biases from DNA re-polymerization when assessing resection progression at this side of the break. The calculated values for each 300 nt fraction were plotted in graphs representing time (h) against distance (kb), enabling the visualization of the bidirectional resection dynamics over time (Fig 2B, bottom graph). This analysis confirmed that the *rad51Δ psy2Δ* strain required more time to reach the *U2* donor sequence compared to the *rad51Δ* mutant, with this delay being specific to the left side of the HO break.

To further analyze resection dynamics, we separated the left and right sides of the graph and use the trendline equations to calculate resection velocities for both sides of the HO break in *rad51Δ* and *rad51Δ psy2Δ* strains (Fig 2C). In the control *rad51Δ* strain, resection rates were 3.52 kb/h and 3.74 kb/h on the left and right sides of the HO break, respectively (Fig 2C). In the *rad51Δ psy2Δ* strain, the resection rate on the left side decreased to 2.73 kb/h (Fig 2C, left graph), confirming a slower resection rate on this side on the break in the absence of Psy2. Interestingly, the right-side resection rate increased slightly to 5.37 kb/h (Fig 2C, right graph), likely due to the extended resection tracts generated on this side as a result of the longer time required for resection to reach the *U2* donor sequence on the left side of the HO break (Fig 2B, top panels). Supporting this hypothesis, the timing of DNA synthesis on the right side of the HO break increased from 12 h in *rad51Δ* cell to 16 h in *rad51Δ psy2Δ* cells (Fig 2D). Furthermore, analysis of the average coverage of a genomic fragment on the right side of the HO break comprised between coordinates 101 kb and 104 kb over a 0-24 h time interval revealed no defects in DNA resection but retrieved a significant delay in the repolymerization of the resected DNA in *rad51Δ psy2Δ* cells compared to *rad51Δ* cells (Fig 2E).

We conclude that SSA repair analysis using a GWS approach provides a highly sensitivity method for detecting subtle variations in resection rates and symmetry at the HO surroundings. Applying this genomic strategy to cells lacking Psy2 revealed that PP4^Psy2^ is essential for an efficient resection to the left side of the HO site, while is dispensable for sustaining resection on the right side. Left-specific stimulation of resection by PP4^Psy2^ ensures timely repair of the HO-induced break by SSA.

### Resection and SSA repair defects of *psy2Δ* cells are alleviated when expressing a phosphodeficient H2A version or in cells lacking Rad9

Previous studies have demonstrated that Pph3 plays a determinant role in dephosphorylating histone H2A following DNA damage to promote efficient resection (Keogh *et al*., 2006). Elevated phosphorylation levels of histone H2A are known to adversely affect DNA resection (Villoria *et al*., 2019). It has also been established that attenuation of Rad53 phosphorylation levels through the expression of a phospho-deficient histone H2A variant suppresses the resection defects of *pph3Δ* cells (Villoria *et al*., 2019). In this context and considering the similarities in resection and DNA repair defects of *pph3Δ* and *psy2Δ* mutants, we wondered whether the expression of a phospho-deficient H2A variant could alleviate *psy2Δ* defects in SSA.

To thoroughly analyze the rate and symmetry of resection in Southern blot experiments, we designed a new set of probes that hybridized at -22 kb, -16 kb, +9 kb and +16 kb from the HO cleavage site (Fig 3A, yellow labeled probes). This set of probes, combined with those previously designed for DNA repair analysis (Fig 1A and Fig 3A, green labeled probes), provides a reliable system to determine symmetry and dynamics of resection. Expression of a phospho-deficient histone H2A variant in which serine 129 has been substituted by a stop codon in both *HTA1* and *HTA2* alleles (*hta1/hta2-S129\**) (Downs *et al*, 2000), significantly improved SSA of *rad51Δ psy2Δ* cells. This improvement is denoted by the faster disappearance of the donor band signal, and consequently, the earlier accumulation of the repair product compared to control *rad51Δ psy2Δ* cells (Fig 3B, top panel). Consequently, the I_D/P_ value decreased from 14.76 h in *rad51Δ psy2Δ* cells to 7.28 h in *rad51Δ psy2Δ hta1/2-S129\** cells (Fig 3C, top graphs). The intensity of all resection band signals diminished more rapidly in the *rad51Δ psy2Δ hta1/2-S129\** strain compared to the *rad51Δ psy2Δ*, indicating enhanced resection progression on both sides of the break (Fig 3B, bottom panel and Fig 3C, bottom graphs). Notably, the delay in the resection profile of the -22 kb band observed in *rad51Δ psy2Δ* cells was abolished in cells expressing the phospho-deficient H2A variant (Fig 3C, bottom graphs), indicating that excessive H2A phosphorylation may account for the asymmetric resection seen in cells lacking Psy2. As a result of the faster resection dynamics in *rad51Δ psy2Δ hta1/2-S129\** cells, DNA re-polymerization kinetics at +16 kb and +9 kb were enhanced compared to *rad51Δ psy2Δ* control cells (Fig 3C, bottom graphs). The overall improvement in HO repair in cells lacking H2A phosphorylation was accompanied by reduced Rad53 phosphorylation levels during the HO induction (Fig 3D, left panel). Finally, absence of Psy2 significantly increased H2A phosphorylation levels in response to HO expression (Fig 3D, right panel), indicating that PP4^Psy2^ avoids H2A hyper-phosphorylation during SSA. In summary, these results confirm that the left-HO resection defects observed in the absence of PP4^Psy2^ activity are bypassed when expressing a non-phosphorylatable version of H2A.

**Figure 3.**
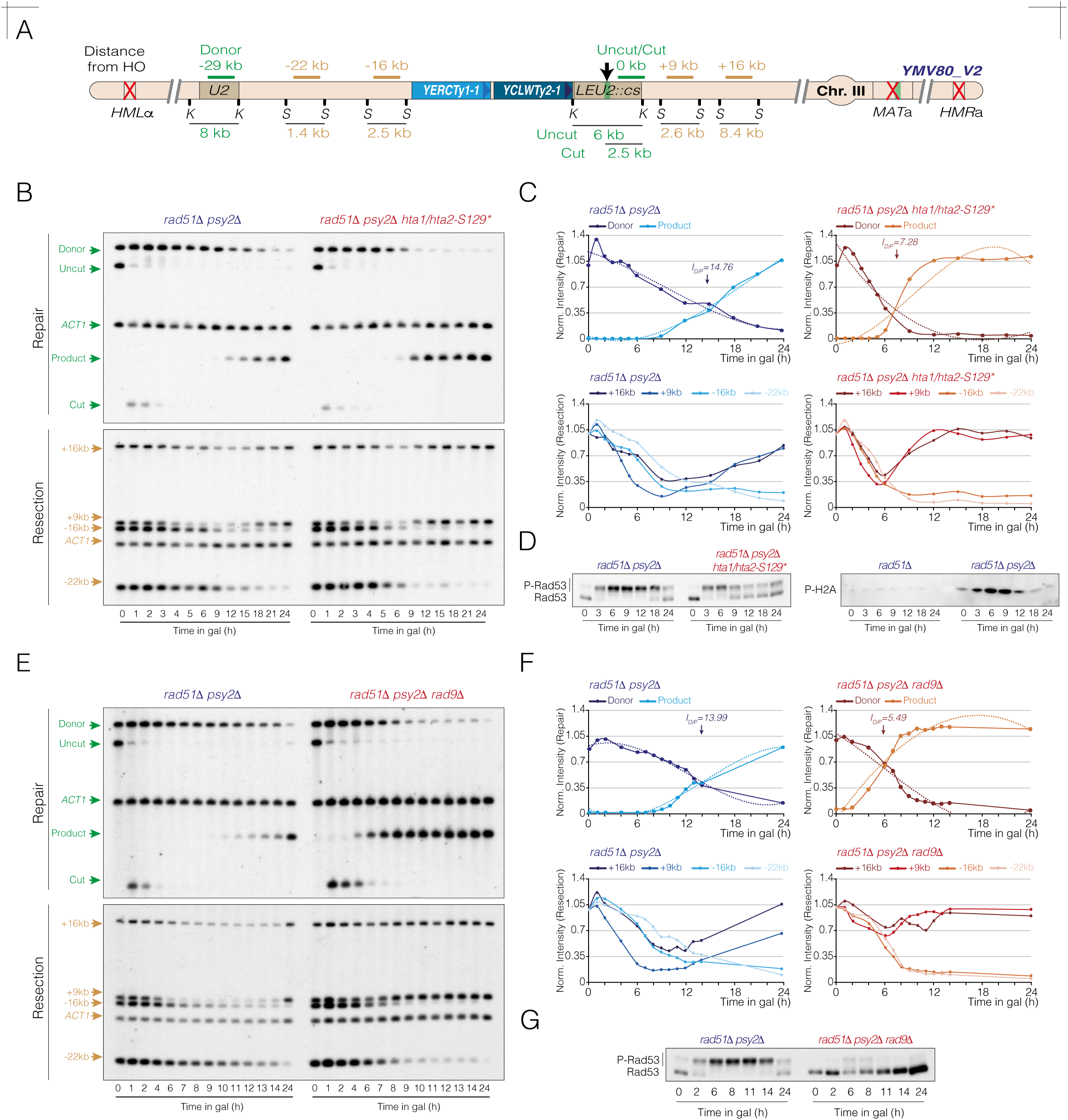
Resection and SSA repair defects of *psy2Δ* cells are mitigated by expressing a phosphor-deficient H2A variant or by deleting *RAD9*. **A.** Schematic representation illustrating the set of probes designed to assess DNA repair and resection efficiency in the *YMV80_v2* strain. The distances from the HO break and the sizes of the DNA fragments generated after digesting with *Kpn*I or *Sty*I are indicated. **B.** Southern blot analysis of *rad51Δ psy2Δ* and *rad51Δ psy2Δ hta1/2-S129\** cells carrying the DNA repair system depicted in A. Cells were grown overnight in YP-Raffinose and the HO nuclease was induced by adding galactose to the media. Samples were taken at various time points, genomic DNA was extracted, digested with *Kpn*I (top panel) or *Sty*I (bottom panel), and analyzed by Southern blot using the probes described in A. The *ACT1* gene sequence was used as loading control. **C.** Top graphs represent the quantification of the donor and product band signals from B (top panel), averaged from two biological replicates and normalized to their respective *ACT1* and uncut T0 signals. The intersection point of the donor and product trend lines (I_D/P_ value) represents the mean from two independent experiments. The statistical significance (*P*-value) of differences between both strains assessed by a two-tailed unpaired Student’s *t-*test is 0.0094 (**). Bottom graphs represent the quantification of +16 kb, +9 kb, -16 kb and -22 kb band signal intensities from B (bottom panel), averaged from two experiments and normalized against their respective *ACT1* and uncut T0 signals. **D.** Samples from the experiment shown in B were collected at the indicated time points, and proteins were extracted using TCA precipitation. Western blot analysis was performed to detect Rad53 phospho-isoforms (left panel) and γ-H2A phosphorylation (right panel). **E.** Southern blot analysis of *rad51Δ psy2Δ* and *rad51Δ psy2Δ rad9Δ* cells carried out under the same experimental conditions as in B to evaluate DNA repair (top panel) and resection (bottom panel) efficiencies. **F.** Top graphs represent quantification of the donor and product band signals from E (top panel), averaged from two biological replicates and normalized to their respective *ACT1* and uncut T0 band signals. The intersection point of the donor and product trend lines (I_D/P_ value) represents the mean from two independent experiments. The statistical significance (*P*-value) of differences between both strains assessed by a two-tailed unpaired Student’s *t-*test is 0.0036 (**). Bottom graphs are obtained from the data shown in E (bottom panel), representing the averaged +16 kb, +9 kb, -16 kb and -22 kb band signal intensities from two independent experiments, normalized against their respective *ACT1* and uncut T0 band signals. **G.** Protein samples at the indicated time points from the experiment shown in E were TCA extracted and subjected to western blotting analysis to assess Rad53 phosphorylation along the HO induction.

It has been documented that phosphorylation of histone H2A stimulates the binding of the adaptor protein Rad9 to the DSB vicinity, which acts as a physical barrier for the Dna2-Sgs1 exonuclease complex, thus affecting DNA resection (Villoria *et al*., 2019). It has also been established that the attenuated Rad53 phosphorylation levels inherent of *rad9Δ* mutants suppress the resection defects of *pph3Δ* cells (Villoria *et al*., 2019). Depletion of *RAD9* in *rad51Δ psy2Δ* cells also had a beneficial effect on SSA efficiency (Fig 3E, top panel and Fig 3F top graphs), resection dynamics (Fig 3E bottom panel and Fig 3F bottom graphs) and Rad53 dephosphorylation (Fig 3G), suggesting that PP4^Psy2^ is required to sustain a steady-state phosphorylation of Rad53 during SSA, thus preventing the negative impact of Rad9 on resection. This molecular mechanism seems to be particularly important to ensure an efficient resection on the left side of the HO break.

### Asymmetric resection of *psy2Δ* cells arise from the presence of retrotransposons elements adjacent to the HO cleavage site

We have seen above that absence of Psy2 particularly affects resection on the left side of the HO cleavage site. As commented earlier, the analysis of the inter-*U2* sequence revealed two tandem Ty retrotransposons elements, *YERCTy1-1* and *YCLWTy2-1*, located adjacent to the cleavage site (Fig 2A). Due to their repetitive nature, these genomic features have the potential to form complex secondary DNA structures that could interfere with transcription, replication or resection (Aguilera & Garcia-Muse, 2012; Ben-Aroya *et al*, 2004; Campos *et al*., 2023; Casper *et al*, 2009; Ramos *et al*., 2022; Zeng *et al*, 2021). Based on this, we hypothesized that if the role of PP4^Psy2^ in regulating Rad53 and H2A phosphorylation in response to a DSB is particularly necessary for enhancing resection through these sequences, then disrupting the Ty elements adjacent to the HO site should bypass its requirement in SSA.

To test this hypothesis, we replaced the entire *YERCTy1-1/YCLWTy2-1* region for a *KanMX-URA3* cassette using the *delitto perfetto* system (Storici & Resnick, 2006) and re-analyzed resection efficiency. Exponentially growing cultures of *rad51Δ psy2Δ* and *rad51Δ psy2Δ yercty1-1/yclwty2-1::KanMX-URA3* were HO induced and samples were collected at various time points to assess resection rate and symmetry by Southern blot (Fig 4A). Deletion of the sequences encoding the two Ty elements significantly improved DNA end resection to the left side of the HO break, as denoted by the rapidly decrease of the donor signal in the *rad51Δ psy2Δ yercty1-1/yclwty2-1::KanMX-URA3* strain compared to the *rad51Δ psy2Δ* control strain (Fig 4B, top panel). This improvement in the resection rate of *rad51Δ psy2Δ yercty1-1/yclwty2-1::KanMX-URA3* cells resulted in a faster accumulation of the DNA repair product, as indicated by the reduced I_D/P_ value obtained when compared to the *rad51Δ psy2Δ* strain (I_D/P_ = 8.46 h and 15.92 h, respectively) (Fig 4C, top graphs).

**Figure 4.**
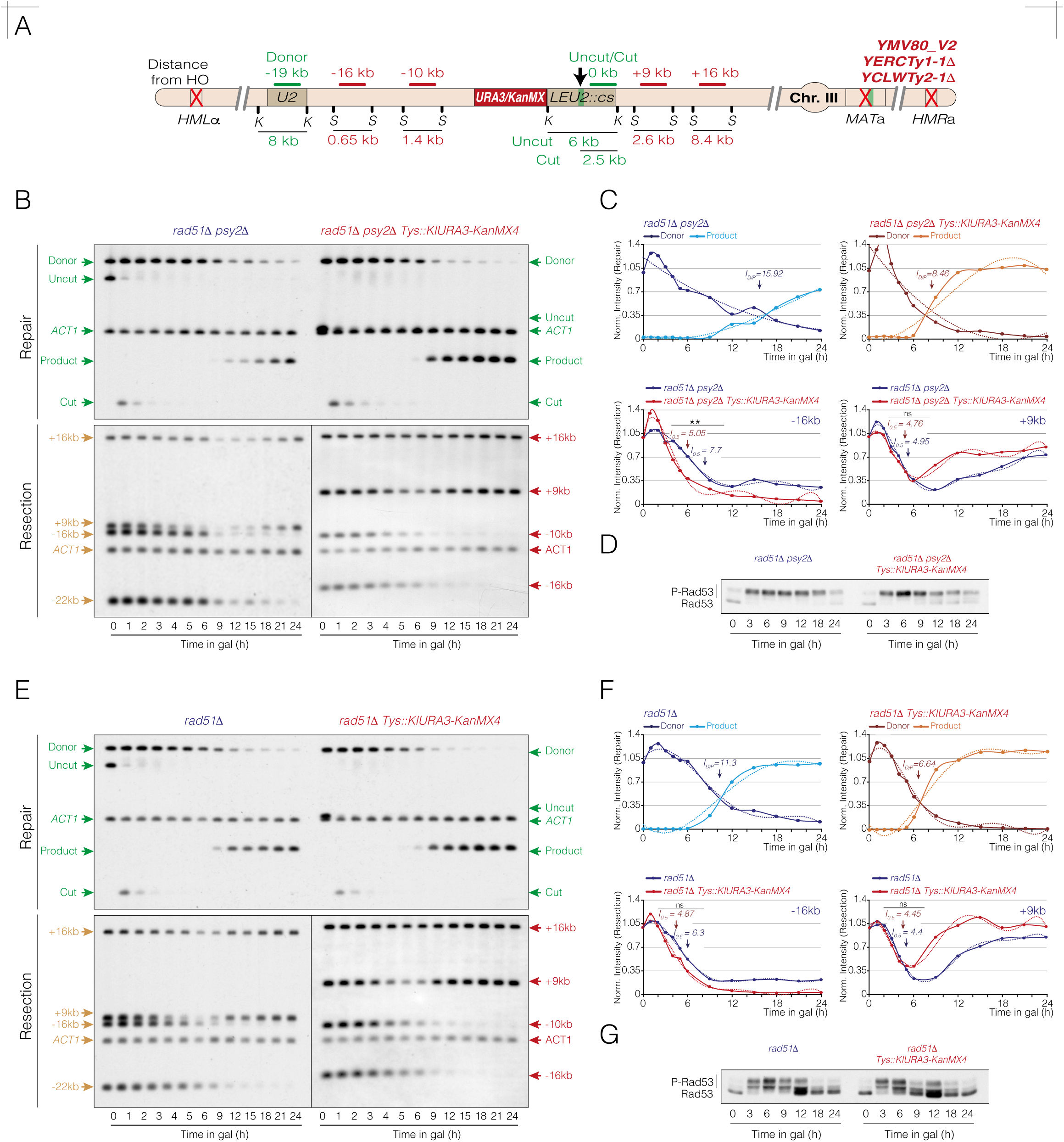
Asymmetric resection in the absence of Psy2 is bypassed after removing the Ty elements located left to the HO cleave site. **A.** Schematic representation showing the set of probes designed to assess DNA repair and resection efficiency in the *YMV80_v2 yercty1-1Δ yclwty2-1Δ* strain. The distances from the HO cleavage site and the sizes of the DNA fragments generated after digestion with *Kpn*I or *Sty*I are indicated. **B.** Southern blot analysis of *rad51Δ psy2Δ* and *rad51Δ psy2Δ yercty1-1Δ yclwty2-1Δ* strains. Cells were cultured overnight in YP-Raffinose, and the HO nuclease was induced by adding galactose to the medium. Samples were collected at various time intervals, and genomic DNA was extracted, digested with *Kpn*I (top panel) or *Sty*I (bottom panel), and analyzed by Southern blot using the probes shown in A. Note that resection probes in both strains retrieve bands shifted relative to one another due to differences in the DNA fragment sizes resulting from deleting the *YERCTy1-1/YCLWTy2-1* tandem sequence. The *ACT1* gene sequence served as a loading control. *Tys::KlURA3-KanMX4* denotes deletion of the tandem *YERCTy1-1/YCLWTy2-1*. **C.** Top graphs show the quantification of donor and product band signals from B (top panel), averaged from two biological replicates and normalized to their respective *ACT1* and uncut T0 signals. The intersection point of the donor and product trend lines (I_D/P_ value) represents the mean from two independent experiments. The statistical significance (*P*-value) of differences between both strains assessed by a two-tailed unpaired Student’s *t-*test is 0.0009 (***). Bottom graphs represent the quantification of +9 kb and -16 kb band signal intensities from B (bottom panel), averaged from two experiments and normalized against their respective *ACT1* and uncut T0 signals. The I_0.5_ value represents the mean from two independent experiments. The statistical significance of differences between both strains has been assessed using a two-tailed unpaired Student’s *t-*test. *Tys::KlURA3-KanMX4* denotes deletion of the tandem *YERCTy1-1/YCLWTy2-1*. **D.** Samples from the experiment shown in B were collected at the indicated time points. Proteins were extracted using TCA precipitation and subjected to western blotting to monitor Rad53 phosphorylation dynamics during HO induction. *Tys::KlURA3-KanMX4* denotes deletion of the tandem *YERCTy1-1/YCLWTy2-1*. **E.** Southern blot analysis of *rad51Δ* and *rad51Δ yercty1-1Δ yclwty2-1Δ* strains following the same experimental condition shown in B. *Tys::KlURA3-KanMX4* denotes deletion of the tandem *YERCTy1-1/YCLWTy2-1*. **F.** Top graphs present the quantification of donor and product band signals from E (top panel), averaged from two biological replicates and normalized to their respective *ACT1* and uncut T0 signals. The intersection point of the donor and product trend lines (I_D/P_ value) represents the mean from two independent experiments. The statistical significance (*P*-value) of differences between both strains assessed by a two-tailed unpaired Student’s *t-*test is 0.005 (**). Bottom graphs represent the quantification of band signal intensities at +9 kb and -16 kb from E (bottom panel), averaged from two experiments and normalized against their respective *ACT1* and uncut T0 signals. The I_0.5_ value represents the mean from two independent experiments. The statistical significance of differences between both strains has been assessed using a two-tailed unpaired Student’s *t-*test. *Tys::KlURA3-KanMX4* denotes deletion of the tandem *YERCTy1-1/YCLWTy2-1*. **G.** Samples from the experiment shown in E were collected at the indicated time points, proteins were TCA-extracted and subjected to western blotting to monitor Rad53 phosphorylation dynamics during HO induction. *Tys::KlURA3-KanMX4* denotes deletion of the tandem *YERCTy1-1/YCLWTy2-1*.

Although replacing *YERCTy1-1* and *YCLWTy2-1* transposable elements with the *KanMX-URA3* cassette partially compensates for the disrupted DNA region, the distance between the two *U2* sequences remains slightly shorter in the *yercty1-1/yclwty2-1::KanMX-URA3* mutant compared to the *rad51Δ psy2Δ* control strain. To overcome this *U2-U2* length discrepancy, we designed a new set of probes specific for the analysis of resection in the *yercty1-1/yclwty2-1::KanMX-URA3* background that perfectly compensate for the inter-*U2* distances measured in the *rad51Δ psy2Δ* control strain (Fig 4A, red probes). Note that the -16 kb probe perfectly match in both *rad51Δ psy2Δ* and *rad51Δ psy2Δ yercty1-1/yclwty2-1::KanMX-URA3* strains (Fig 3A, yellow probes and Fig 4A, red probes), ensuring an accurate comparison of resection dynamics at equidistant distances from the cut site. We repeated the experiment under the same experimental conditions described previously and performed a Southern blot analysis (Fig 4B, bottom panel) using the resection probes depicted in figure 3A and 4A. Quantification of resection dynamics at -16 kb revealed a significant improvement in the *rad51Δ psy2Δ yercty1-1/yclwty2-1::KanMX-URA3* strain compared to *rad51Δ psy2Δ* cells (Fig 4B, bottom panel). The averaged time needed for the cell population to resect the -16 kb (I_0.5_, time needed for a band signal intensity to drop from 1 to 0.5) was 7.7 h in *rad51Δ psy2Δ* cells, whereas this value decreased to 5.05 h in the *rad51Δ psy2Δ yercty1-1/yclwty2-1::KanMX-URA3* strain (Fig 4C, bottom left graph). As expected, the I_0.5_ values on the right side of the HO break (+9 kb) were similar for both strains (Fig 4C, bottom right graph; *rad51Δ psy2Δ* = 4.95 h; *rad51Δ psy2Δ yercty1-1/yclwty2-1::KanMX-URA3* strain = 4.76 h). Disrupting the tandem *YERCTy1-1/YCLWTy2-1* elements in *rad51Δ psy2Δ* cells improved Rad53 dephosphorylation during the latest stages of the damage response (Fig 4D), probably due to the improved SSA repair in these cells. This suggests that the hyper-phosphorylation of Rad53 typically seen in cells lacking Psy2 is not solely attributable to its role in checkpoint silencing after repair, but for their incapacity to timely sustain DNA repair.

To completely rule out that the enhanced left-side resection observed upon disrupting the *YERCTy1-1/YCLWTy2-1* elements was exclusively due to the reduced inter-*U2* distance, we repeated the same experiment in *PSY2*^+^ cells. As anticipated, the *rad51Δ yercty1-1/yclwty2-1::KanMX-URA3* strain exhibited a lower I_D/P_ value (6.64 h) compared to *rad51Δ* control cells (11.3 h), attributable to the shortener *U2-U2* distance (Fig 4E, top panel and Fig 4F, top graphs). Accordingly, the I_0.5_ value on the left side of the HO decreased from 6.3 h to 4.7 h (Fig 4F, bottom left graph). Importantly, this reduction in the I_0.5_ value when removing the tandem Ty elements was less pronounced than that observed in *psy2Δ* backgrounds (Fig 4C bottom left graph), indicating that the presence of these repetitive regions particularly affects cells lacking PP4^Psy2^ activity. As expected, the I_0.5_ values on the right side of the HO break (+9 kb) were similar for both strains (Fig 4F, bottom right graph; *rad51Δ* = 4.4 h; *rad51Δ yercty1-1/yclwty2-1::KanMX-URA3* strain = 4.45 h). Disruption of the tandem *YERCTy1-1/YCLWTy2-1* elements in *rad51Δ* cells slightly improved Rad53 dephosphorylation dynamics during the latest stages of the damage response (Fig 4G), likely due to the enhanced DNA re-synthesis capacity (Fig 4F bottom right graph) resulting from the shortened inter-*U2* distance.

Collectively, these results demonstrate that PP4^Psy2^ in necessary for facilitating resection progression throughout heterochromatic DNA regions containing retrotransposons sequences.

### PP4^Psy2^ is predominantly required to sustain resection throughout tandem retrotransposons elements

During this study, we identified a *rad51Δ psy2Δ* mutant with a shorter uncut band compared to the control strain in Southern blot experiments (Fig 5B). *De novo* long-read genome assembly of this *YMV80* variant (designated as *YMV80_v3* afterwards) revealed a recombination event between the *YERCTy1-1* and *YCLWTy2-1* Ty elements, resulting in the generation of a single retrotransposon composed of the N-terminal region of *YERCTy1-1* and the C-terminal region of *YCLWTy2-1* (Fig 5A). Interestingly, this new *rad51Δ psy2Δ* background (hereafter referred as to *rad51Δ psy2Δ^YMV80_v3^*) did not exhibit drastic defects in DNA repair in Southern blot experiments when compared to the *rad51Δ^YMV80_2^* control strain, neither in the dynamics of the donor band disappearance due to resection, nor in the accumulation of the repair product (Fig 5B). Accordingly, only a slight decrease in the I_D/P_ values was detected in the *rad51Δ psy2Δ^YMV80_v3^* compared to the *rad51Δ ^YMV80_2^* (I_D/P_ values = 8.92 and 9.99 h, respectively) (Fig 5C). Importantly, although Rad53 was hyper-activated in *rad51Δ psy2Δ^YMV80_v3^* during the initial stages of the response, we noticed a notable reduction at later time points compared to the *rad51Δ psy2Δ ^YMV80_v2^* strain (Fig 5D and Fig 1G, right panels). This result reinforces the notion that the innate hyper-phosphorylation of Rad53 in *psy2Δ* cells is not solely linked to its role in checkpoint inactivation after repair, but for its incapacity to support DNA repair in a timely manner.

**Figure 5.**
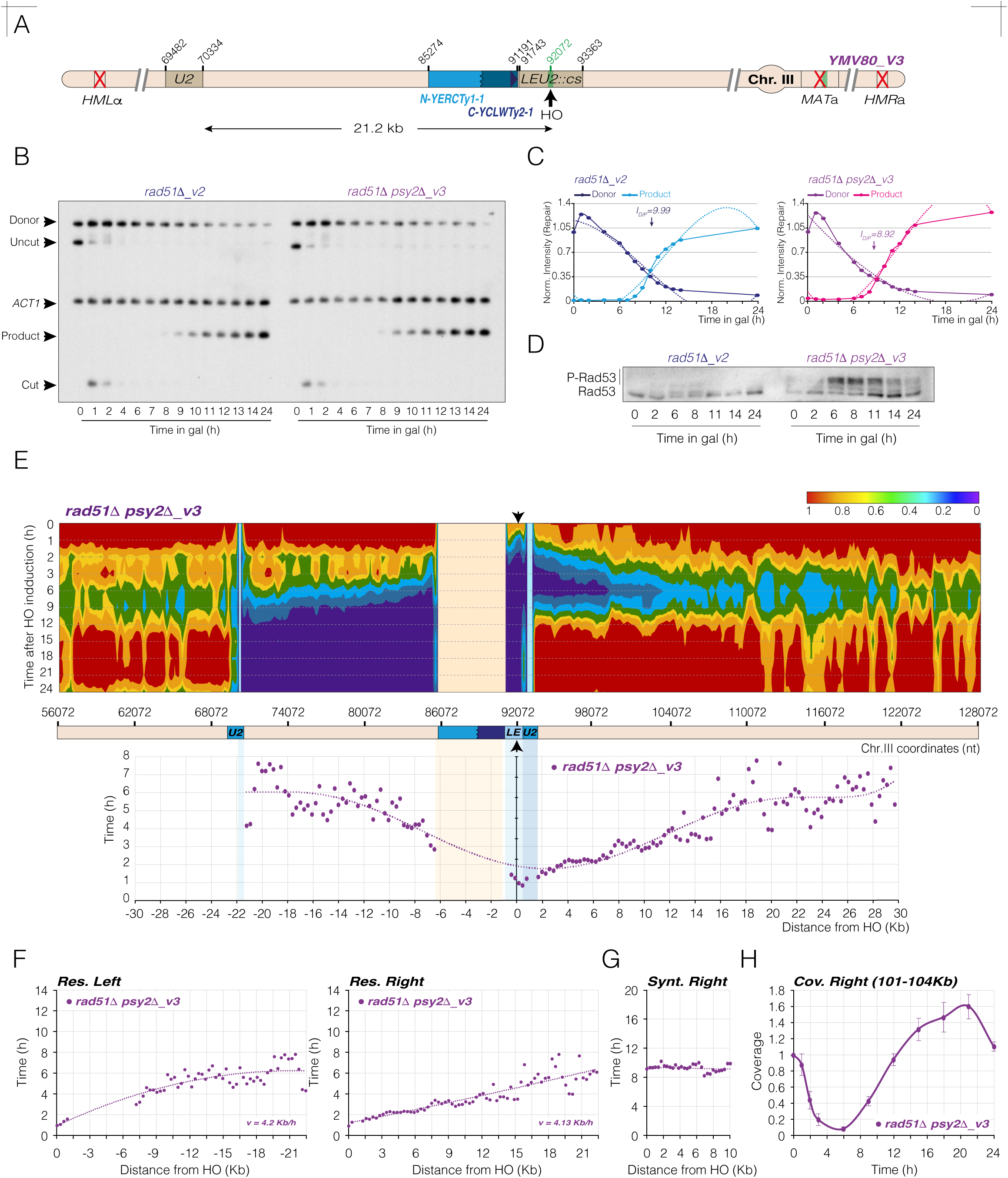
PP4^Psy2^ enhances resection through tandem retrotransposons elements. **A.** Schematic representation illustrating the distinctive genomic features of the *YMV80_v3* strain, as determined by long-read sequencing assembly. The coordinates for each feature and the inter-*U2* distance are shown. **B.** Southern blot analysis of *rad51Δ_v2* and *rad51Δ psy2Δ* _*v3* strains. Cells were grown overnight in YP-Raffinose, and the HO nuclease was induced by adding galactose to the medium. Samples were collected at different time intervals, and genomic DNA was extracted, digested with *Kpn*I, and analyzed by Southern blot using the probes shown in figure 1A. The *ACT1* gene sequence was used as a loading control. **C.** Graphs show the quantification of the donor and product band signals from B, averaged from two biological replicates and normalized to their respective *ACT1* and uncut T0 signals. The intersection point of the donor and product trend lines (I_D/P_ value) represents the mean from two independent experiments. The statistical significance (*P*-value) of differences between both strains assessed by a two-tailed unpaired Student’s *t-*test is 0.604 (ns). **D.** Samples from the experiment shown in B were collected at the indicated time points. Proteins were extracted using TCA precipitation and subjected to western blotting to monitor Rad53 phosphorylation dynamics during HO induction. **E.** Colormap charts displaying coverage levels of the *rad51Δ psy2ΔΔv3* strain. Coverage levels were normalized against the T0 sample and represented by red, green and blue gradients. The *x-*axis and *y-*axis correspond to chromosome III coordinates and time after HO induction, respectively. Black arrow indicates the position of the HO cleavage site. Graph represents the mean of two biological experiments. Note that the Ty element and *U2* sequences (red and blue areas, respectively) were excluded from the analysis. Bellow, a graph representing time (h) *versus* distance (kb) across a +/-30 kb region from the HO site. This was calculated by analyzing the trend line equations for 300 nt average coverage fractions as they dropped from 1 to 0.5 (see Materials and Methods for details). **F.** Analysis of resection dynamics in *rad51Δ psy2ΔΔ v3* cells. The left and right HO sides from the graph depicted in E were split and analyzed separately. Trend line equations were derived from the time (*y*-values) required for resection to cover specific distances (*x*-values) up to 22 kb at both sides of the HO break. Resection rates (*v*) were calculated by measuring the time (*Δt*) needed for resection to accomplish this distance (*Δs*) at both sides of the break (see Materials and Methods for details). **G.** Analysis of DNA re-synthesis in a *rad51Δ psy2ΔΔv3* strain. Synthesis rate was calculated by averaging coverage levels of 300 nt fractions spanning from 0 kb to 10 kb to the right of the HO site during the 9 h – 24 h time interval. The trend line equations for each fraction were used to determine the time (h) required for coverage values to recover to 0.5 (see Materials and Methods for details). **H.** Analysis of coverage levels on the right side of the HO site in a *rad51Δ psy2ΔΔv3* strain after inducing the HO nuclease. The averaged coverage of a right HO region comprised between coordinates 101,000 and 104,000 on chromosome III was calculated and plotted over time.

It is worth mentioning that in the *YMV80_v3* background, the distance between the HO break and the donor *U2* sequence is shortened to 21.2 kb (Fig 5A). To ensure that this reduced inter-*U2* distance does not solely account for the enhanced repair observed in *rad51Δ psy2Δ ^YMV80_v3^* cells, we applied the genomic approach described earlier (Fig 2) to analyze the kinetics and symmetry of resection in this strain. HO expression was induced in exponentially growing cells, and samples were collected at distinctive time intervals for genome-wide sequencing. The sequenced reads were aligned to the *YMV80_v3* reference genome, and coverage levels were normalized to the undamaged T0 sample to generate 2D (Fig EV3A), 3D (Fig EV3B) and heatmap (Fig 5E) coverage profiles. The heatmap coverage profile for the *rad51Δ psy2Δ ^YMV80_v3^* mutant showed a gradual and symmetrical loss of coverage on both sides of the break during the 1-9 h time interval following HO induction (Fig 5E, top panel, Fig EV3A-B). Validating Southern blot experiments, inter-*U2* resection in the *rad51Δ psy2Δ^YMV80_v3^* was more efficient than in the *rad51Δ psy2Δ ^YMV80_v2^*, as denoted by the rapid transition from red to blue coverage values within the *U2*-*U2* region (Fig 5E and Fig 2B, bottom panel). Accordingly, resection in *rad51Δ psy2Δ ^YMV80_v3^* cells reached the *U2* donor sequence by 9 h after HO induction, a time point at which coverage levels on the right side of the HO break began to recover. These findings suggest that both resection dynamics and repair efficiency are improved in the *rad51Δ psy2Δ^YMV80_v3^* strain compared to the *rad51Δ psy2Δ^YMV80_v2^* strain.

Quantification of the resection rates on both sides of the HO break in the *rad51Δ psy2Δ^YMV80_v3^* strain (Fig 5E, bottom graph) confirmed that resection progressed at similar rates on both sides of the break (Fig 5F), with calculated velocities of V_left_ = 4.2 kb/h and V_right_ = 4.13 kb/h. It is important to note that, while the overall resection rates between 0 kb and 21 kb were comparable on both sides of the break, the initial resection on the left side was slightly restrained as it traversed the Ty sequence. This lag was later compensated by increasing the resection rate form 12 kb to 21 kb (Fig 5F). This effect, also observed in the *rad51Δ YMV80_v2* strain, supports the notion that, although to a lesser extent, single Ty elements influence resection progression. Still, and in accordance with the similar net resection rate observed at both sides of the break, DNA synthesis to the right of the HO break in the *rad51Δ psy2Δ^YMV80_v3^* occurred faster than in *rad51Δ^YMV80_v2^* cells at approximately 9 h post-HO induction (Fig 5G). Furthermore, the average coverage between coordinates 101-104 kb over the 0-24 h time interval of the *rad51Δ psy2Δ^YMV80_v3^* strain (Fig 5H) closely matched that of the *rad51Δ^YMV80_v2^* strain (Fig 2E), confirming an efficient and timely DNA repolymerization of the resected DNA.

These results indicate that PP4^Psy2^ is required for efficient resection through regions enriched with retrotransposon element, particularly when these elements are arranged as tandem repeats. Given the repetitive nature of these sequences, these results anticipate that in the absence of PP4^Psy2^ activity, unresolved DNA secondary structures might form ahead of the resection machinery, impeding its progression.

### Pif1 and Rrm3 helicases exhibit PP4^Psy2^ -dependent synergistic roles in facilitating resection through tandem retrotransposons

We have shown above that lack of PP4^Psy2^ activity affects resection through tandem Ty sequences, likely due to the existence of unresolved secondary DNA structures obstructing resection progression. It has been postulated that Rad53 phosphorylates and regulates Rrm3 and Pif1 helicases during replication stress (Rossi *et al*, 2015). Since the absence of Psy2 leads to a hyper-phosphorylation state of Rad53 in response to an HO break (Fig 1G), it is plausible that the lack of PP4^Psy2^ might influence Pif1 and Rrm3 phosphorylation dynamics during SSA.

To test this hypothesis, we analyzed the phosphorylation profile of Pif1 and Rrm3 during the induction of an HO break in the *YMV80_v2* background in wild type and *psy2Δ* cells. Two hours from the HO induction resulted in the accumulation of high molecular weight bands of Pif1 and Rrm3 in both strains (Fig 6A,B), indicating that both proteins are phosphorylated in response to the HO cut in a Psy2-independent manner. However, while phosphorylation of Pif1 and Rrm3 was transient in wild type cells, it remained stabilized through the entire experiment in the absence of *PSY2* (Fig 6A,B). This result suggests that PP4^Psy2^ is required to endorse Pif1 and Rrm3 dephosphorylation during SSA.

**Figure 6.**
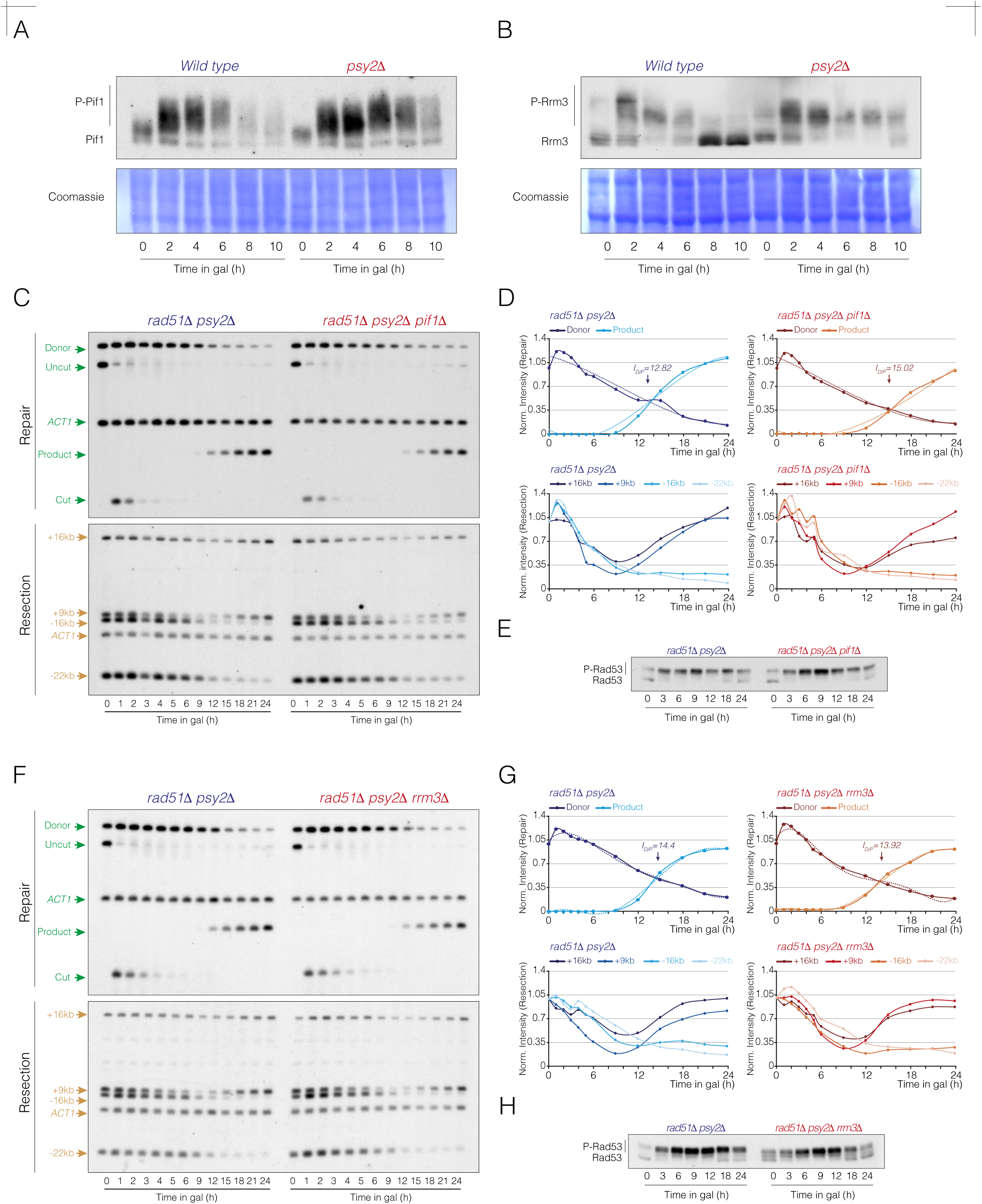
PP4^Psy2^-dependent regulation of Pif1 and Rrm3 helicases is required for DNA repair by SSA. **A.** Western blot of wild type and *psy2Δ* cells to assess the Pif1 phosphorylation profile in SSA. Cells were grown overnight in YP-Raffinose, and the HO nuclease was induced by adding galactose to the medium. Samples were collected at various time points, and proteins were extracted using TCA precipitation and subjected to Phos-tag western blotting to monitor Pif1 phosphorylation dynamics. Coomassie staining is shown as loading control. **B.** Western blot analysis of wild type and *psy2Δ* strains to assess Rrm3 phosphorylation in SSA following the same experimental condition as in A. Coomassie staining is shown as loading control. **C.** Southern blot analysis of *rad51Δ psy2Δ* and *rad51Δ psy2Δ pif1Δ* strains. Cells were grown overnight in YP-Raffinose before adding galactose to induce HO expression. Samples were taken at different time points, and genomic DNA was extracted, digested with *Kpn*I (top panel) or *Sty*I (bottom panel), and analyzed by Southern blot using the probes illustrated in figure 3A. The *ACT1* gene sequence was used as loading control. **D.** Top graphs show the quantification of donor and product band signals in C (top panel), averaged from two biological replicates and normalized to their respective *ACT1* and uncut T0 signals. The intersection point of the donor and product trend lines (I_D/P_ value) represents the mean from two independent experiments. The statistical significance (*P*-value) of differences between both strains assessed by a two-tailed unpaired Student’s *t-*test is 0.109 (ns). Bottom graphs represent the quantification of +16 kb, +9 kb, -16 kb and -22 kb band signal intensities from C (bottom panel), averaged from two independent experiments and normalized against their respective *ACT1* and uncut T0 signals. **E.** Samples from the experiment shown in C were collected at the indicated time points. Proteins were extracted using TCA precipitation and subjected to western blotting to monitor Rad53 phosphorylation in response to the HO induction. **F.** Southern blot analysis of *rad51Δ psy2Δ* and *rad51Δ psy2Δ rrm3Δ* strains following the same experimental condition as in C. Samples were taken at distinctive time intervals, and genomic DNA was extracted, digested with *Kpn*I (top panel) or *Sty*I (bottom panel), and analyzed by Southern blot using the probes illustrated in figure 3A. The *ACT1* gene sequence was used as loading control. **G.** Top graphs display the quantification of donor and product band signals from F (top panel), averaged from two biological replicates and normalized to their respective *ACT1* and uncut T0 signals. The intersection point of the donor and product trend lines (I_D/P_ value) represents the mean from two independent experiments. The statistical significance (*P*-value) of differences between both strains assessed by a two-tailed unpaired Student’s *t-*test is 0.4485 (ns). Bottom graphs represent the quantification of +16 kb, +9 kb, -16 kb and -22 kb band signal intensities from C (bottom panel), averaged from two independent experiments and normalized to their respective *ACT1* and uncut T0 signals. **H.** Samples from the experiment shown in F were taken at the indicated time points, proteins were TCA extracted and subjected to western blotting to assess the Rad53 phosphorylation profile in response to HO induction.

Next, we evaluate SSA repair efficiency in *rad51Δ* cells combined with *pif1Δ*, *rrm3Δ* and *pif1Δ rrm3Δ*. Disruption of *PIF1* resulted in minor defect in SSA repair, both in terms of donor disappearance and product formation (Fig EV4A, top panel). The calculated I_D/P_ values for *rad51Δ* and *rad51Δ pif1Δ* cells were 10.72 h and 11.67 h, respectively (Fig EV4B, top graphs). Consistently, resection dynamics (Fig EV4A, bottom panel and Fig EV4B, bottom graphs) and Rad53 phosphorylation profiles (Fig EV4C) were comparable between *rad51Δ* and *rad51Δ pif1Δ* cells. We neither retrieved drastic changes when disrupting *RRM3* in SSA efficiency, either in the repair of the HO break (I_D/P_ *rad51Δ* = 10.96 h; I_D/P_ *rad51Δ rrm3Δ* = 11.47 h) (Fig EV4D, top panel and Fig EV4E, top graphs), resection (Fig EV4D, bottom panel and Fig EV4E, bottom graphs) or Rad53 phosphorylation (Fig EV4F). However, the combination of *PIF1* and *RRM3* depletions in *rad51Δ* cells drastically restrained SSA repair (Fig EV4G, top panel). This was reflected in a significant reduction in the I_D/P_ value from 10.25 h in *rad51Δ* cells to 14.3 h in the *rad51Δ pif1Δ rrm3Δ* mutant (Fig EV4H, top graphs). We observed a general delay in resection, accompanied by a slower re-accumulation of the +9 kb and +16 kb band signals (Fig EV4G, bottom panel and Fig EV4H, bottom graphs). Consistent with the reduced levels of resection, we detected a modest Rad53 activation in *rad51Δ pif1Δ rrm3Δ* mutant compared to *rad51Δ* cells (Fig EV4I). These results suggest that Pif1 and Rrm3 helicases cooperate during SSA by promoting efficient DNA end resection in the YMV80 background, with their roles becoming particularly critical in the absence of redundancy.

To determine whether PP4^Psy2^ acts over Pif1 or Rrm3, we checked SSA efficiency in *rad51Δ psy2Δ* cells combined with single *pif1Δ* and *rrm3Δ* mutants. The DNA repair defects observed in the *rad51Δ psy2Δ* mutant were exacerbated in *rad51Δ psy2Δ pif1Δ* cells, as reflected by impaired formation of the product band (Fig 6C, top panel). Accordingly, the I_D/P_ value increased from 12.82 h in *rad51Δ psy2Δ* to 15.02 h in *rad51Δ psy2Δ pif1Δ* cells (Fig 6D, top graphs). Delayed I_D/P_ value was attributed to a failure in the accumulation of the repair product, likely due to the role of Pif1 in stimulating DNA synthesis (Buzovetsky *et al*, 2017). Importantly, this defect was independent of resection dynamics, as both strains displayed comparable kinetics in the donor signal loss (Fig 6C, top panel and Fig 6D, top graph) and similar resection bands profiles (Fig 6C, bottom panel and Fig 6D, bottom graph). Moreover, Rad53 phosphorylation profiles did not differ significantly between *rad51Δ psy2Δ* and *rad51Δ psy2Δ pif1Δ* cells (Fig 6E). We neither detected additional defects when disrupting *RRM3* in *rad51Δ psy2Δ* cells, either in SSA repair (Fig 6F, top panel and Fig 6G, top graphs), resection progression (Fig 6F, bottom panel and Fig 6G, bottom graphs) or Rad53 phosphorylation (Fig 6H).

Overall, these data suggest that Pif1, Rrm3 and PP4^Psy2^ share overlapping functions in facilitating SSA by operating through a common pathway to stimulate resection processivity through tandem Ty sequences.

### Overexpression of Pif1 affects long-range resection on both sides of the DNA break, whereas overexpression of Rrm3 compensate for the loss of Psy2 in SSA

We have seen above that absence of Psy2 leads to a persistent Pif1 phosphorylation during the response to an HO break (Fig 6A), suggesting that PP4^Psy2^ might modulate its helicase activity during SSA. Based on this observation, we speculated that if the resection defects of *psy2Δ* cells are linked to Pif1 dysregulation, its overexpression might bypass the requirement of PP4^Psy2^ in SSA. It is important to remark that Pif1 overexpression has been associated with cell lethality due to interference with DNA replication (Chang *et al*, 2009; Yoshikawa *et al*, 2011). To test this, we fused a nuclear version of Pif1 (Pif1-m1) under the galactose promoter in *rad51Δ psy2Δ* cells and concomitantly overexpress it with the HO nuclease to assess SSA performance. Pif1-m1 overexpression led to a severe defect in DNA resection, as denoted for the almost completely lack of donor band signal decay in Southern blot experiments (Fig 7A, top panel and Fig 7B, top graphs). Consequently, the formation of the product band was totally abolished (Fig 7A, top panel and Fig 7B, top graphs). Resection progression was drastically affected at both sides of the break, only detecting a slight reduction in the +9 kb band signal after 16 h from the HO induction (Fig 7A, bottom panel and 7B, bottom graphs). Accordingly, Rad53 phosphorylation was inefficient, as denoted by the persistent presence of unphosphorylated isoforms throughout the experiment (Fig 7C). These findings indicate that elevated levels of Pif1 are incompatible with DNA end resection, thereby preventing DNA repair by SSA.

**Figure 7.**
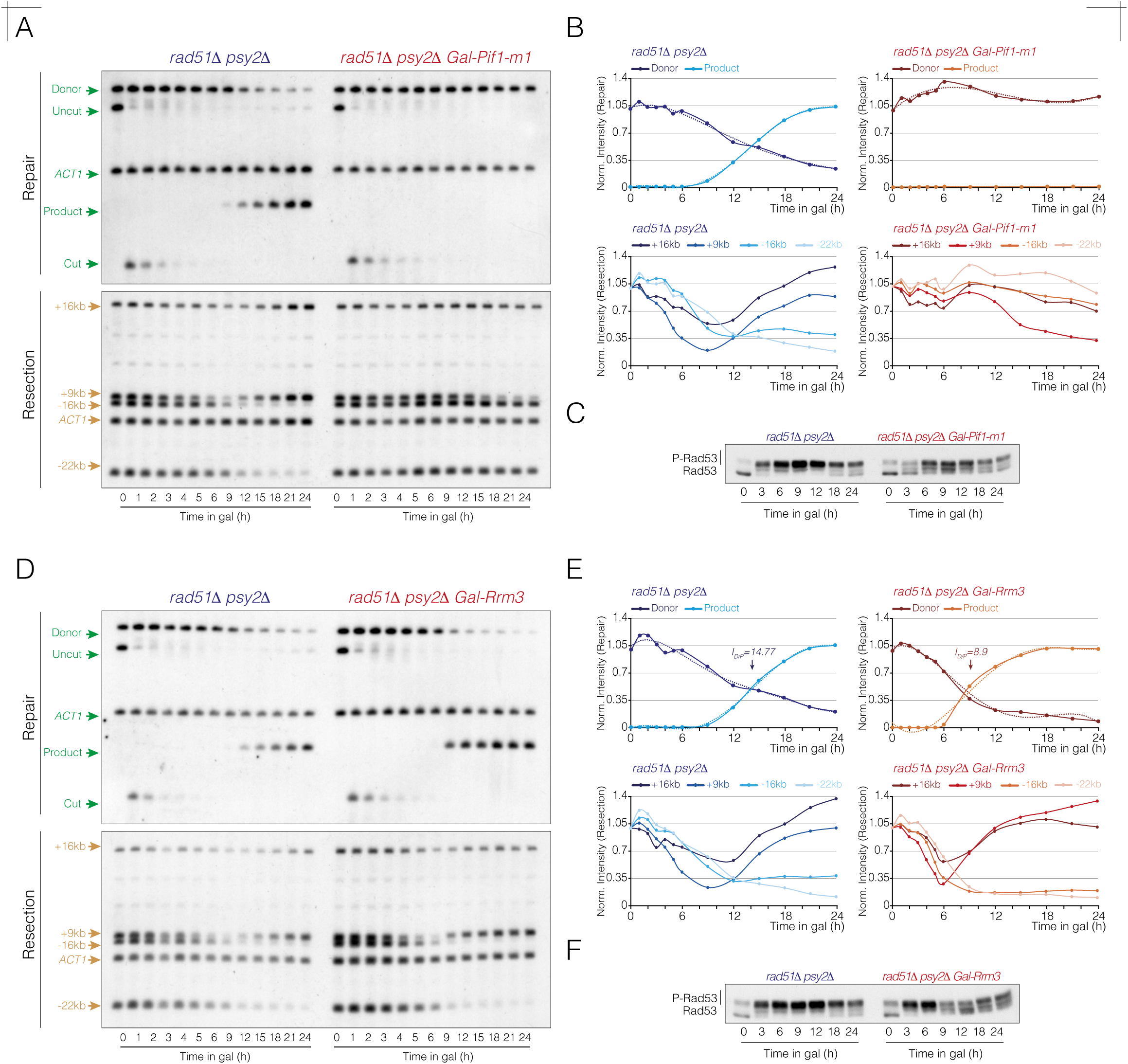
Overexpression of Rrm3, but not Pif1, bypasses Psy2 requirement in SSA. **A.** Southern blot analysis of *rad51Δ psy2Δ* and *rad51Δ psy2Δ GAL-PIF1^m1^* strains. Cells were cultured overnight in YP-Raffinose, and HO and *PIF1^m1^* expression was induced by adding galactose. Samples were collected at various time points, genomic DNA was extracted, digested with *Kpn*I (top panel) or *Sty*I (bottom panel), and analyzed by Southern blot using the probes shown in figure 3A. The *ACT1* gene sequence was used as loading control. **B.** Top graphs depict the quantification of donor and product band signals from A (top panel), averaged from two biological replicates and normalized to their respective *ACT1* and uncut T0 signals. Bottom graphs represent the quantification of +16 kb, +9 kb, -16 kb and -22 kb band signal intensities from A (bottom panel), averaged from two independent experiments and normalized to their respective *ACT1* and uncut T0 signals. **C.** Samples from the experiment shown in A were collected at the indicated time points, proteins were TCA extracted and subjected to western blotting to analyze Rad53 phosphorylation along the response to the HO break. **D.** Southern blot analysis of *rad51Δ psy2Δ* and *rad51Δ psy2Δ GAL-RRM3* strains. Cells were grown overnight in YP-Raffinose before expressing the HO and *RRM3* by adding galactose to the medium. Samples were collected at distinctive time intervals, genomic DNA was extracted, digested with *Kpn*I (top panel) or *Sty*I (bottom panel), and analyzed by Southern blot using the probes shown in figure 3A. The *ACT1* gene sequence was used as loading control. **E.** Top graphs depict the quantification of donor and product band signals from D (top panel), averaged from two biological replicates and normalized to their respective *ACT1* and uncut T0 signals. The intersection point of the donor and product trend lines (I_D/P_ value) represents the mean from two independent experiments. The statistical significance (*P*-value) of differences between both strains assessed by a two-tailed unpaired Student’s *t-*test is 0.002 (**). Bottom graphs represent the quantification of +16 kb, +9 kb, -16 kb and -22 kb band signal intensities from D (bottom panel), averaged from two independent experiments and normalized to their respective *ACT1* and uncut T0 signals. **F.** Samples from the experiment shown in D were collected at the indicated time points. Proteins were TCA extracted and subjected to western blotting to determine Rad53 phosphorylation kinetics along the response to the HO break.

We have also shown that absence of *PSY2* impacts the phosphorylation state of Rrm3 during SSA (Fig 6B), suggesting that PP4^Psy2^ regulates Rrm3 activity. Moreover, we have demonstrated that Psy2 and Rrm3 are epistatic for SSA, indicating that both factors operate within the same pathway to promote efficient resection through transposable repetitive genomic regions. To explore whether Rrm3 overexpression could compensate for the defects in DNA end resection and SSA observed in *psy2Δ* cells, we placed the *RRM3* coding sequence under the control of the galactose promoter and co-express it with the HO nuclease. Overexpression of Rrm3 improved the resection dynamics of *rad51Δ psy2Δ* cells, as evidenced by a faster disappearance of the donor band and an earlier accumulation of the product signal (Fig 7D, top panel). The I_D/P_ values decreased from 14.77 h in the *rad51Δ psy2Δ* strain to 8.9 h under Rrm3 overexpression (Fig 7E, top graphs). We also noted a more rapid loss of the -16 kb and -22 kb band signals, along with an earlier re-accumulation of the +9 kb and +16 kb bands, following resection in *rad51Δ psy2Δ* cells when Rrm3 was overexpressed (Fig 7D, bottom panel and 7E, bottom graphs). Consistently, Rad53 dephosphorylation occurred earlier from the HO induction (Fig 7F). These results demonstrate that Rrm3 overexpression rescues the resection defects of *psy2Δ* cells during SSA and indicate that PP4^Psy2^ modulates Rrm3 activity to ensure efficient resection across tandem Ty-enriched genomic regions.

## Discussion

Phosphorylation in response to DNA damage has been a central focus in the field of DNA repair, particularly on the research of protein kinases involved in the DDR. Although it has been widely accepted that dephosphorylation is also crucial for reversing the effects imposed by the DDR kinases, the specific role and regulation of protein phosphatases in orchestrating the response remains less well documented. To date, much of the research on DDR-dependent protein dephosphorylation has highlighted their role in inactivating the DNA damage checkpoint and in facilitating cell cycle re-entry after DNA repair. However, given the complexity of the DDR pathways and the diverse events performed on each DNA repair stage, it is intuitive to think that a precise phospho-regulation on each step of the repair process might be vital for successfully restore the integrity of the DNA molecule. Supporting this notion, numerous DNA repair factors contain multiple phosphorylation sites that undergo concurrent and dynamic phosphorylation and dephosphorylation, likely involving coordinated action of kinases and phosphatases working in tandem. Importantly, recent studies have underscored the significance of phosphatases in the DDR, revealing their involvement not only as checkpoint recovery factors, but also as factors that directly influence DNA repair processes (Campos & Clemente-Blanco, 2020; Ramos *et al*., 2019). This highlights the dual role of protein phosphatases in fine-tuning the DDR signaling, suggesting a broader scope of their contribution to genome stability.

Here we show that the PP4 phosphatase regulatory element Psy2, but not Psy4, is required for DNA repair by SSA. The absence of Psy2 phenocopies the effects observed in a null catalytic Pph3 mutant, both in DNA repair and Rad53 phosphorylation, confirming its involvement in promoting SSA. In contrast, the absence of Psy4 produces the opposite effect, arguing that Psy4 functions as a repressor of SSA. This observation implies that PP4^Psy2^ and PP4^Psy4^ present antagonist functions in SSA, and that both subunits might compete for regulating Pph3 activity during the DNA damage response. The functional similarities of *pph3Δ* and *psy2Δ* cells in SSA argues for a PP4^Psy2^ role in regulating DNA resection by modulating the phosphorylation status of the checkpoint effector protein Rad53 (Villoria *et al*., 2019). These findings are consistent with previous studies showing that Pph3 preferentially dephosphorylates Rad53 auto-phosphorylation sites, thereby modulating its own activation (O’Neill *et al*., 2007; Villoria *et al*., 2019). In addition to this regulation, absence of *PPH3* or *PSY2* suppresses *mec1-100* lethality upon HO treatment, indicating that PP4^Psy2^ might directly dephosphorylates upstream targets, such as Mec1 (Hustedt *et al*., 2015). Beyond Mec1 and Rad53, H2A is another well documented PP4 substrate in response to MMS and CPT (Chowdhury *et al*., 2008; Keogh *et al*., 2006). Given the elevated levels of phospho-H2A observed in the absence of Psy2 during SSA, it is tempting to speculate that the ultimate role of PP4^Psy2^ in promoting resection may involve chromatin regulation. In this regard, PP4^Psy2^ attenuation of Rad53 phosphorylation prevents excessive accumulation of phospho-Rad9, reducing its chromatin binding capacity, thus enhancing resection progression by the MRX (Ferrari *et al*, 2015) and Dna2-Sgs1 complexes (Bonetti *et al*, 2015; Gobbini *et al*, 2015). Importantly, phosphorylated H2A stimulates Rad9 binding at break sites (Hammet *et al*, 2007; Toh *et al*, 2006), indicating that PP4^Psy2^ might stimulate resection by reducing the presence of Rad9 at the HO surroundings, as previously proposed for Pph3 (Villoria *et al*., 2019). This idea fits with the beneficial effect of removing serine 129 from H2A or depleting Rad9 on SSA performance in *psy2Δ* cells. Importantly, Cdk-dependent phosphorylation of Psy2 is important for an efficient SSA in a Rad53-independent manner, suggesting that in addition to Rad53, PP4^Psy2^ might have additional targets involved in DNA repair by SSA.

Besides the biochemical mechanisms by which PP4^Psy2^ exerts its function in stimulating resection, we have shown that PP4^Psy2^ is required to sustain resection symmetry at both sites of the break. This result infers the existence of “difficult to resect” regions that need a complete functional PP4^Psy2^ complex for its processing. By applying *de novo* long-reads assembly of the *YMV80* background used in this work (Vaze *et al*, 2002) we have detected the presence of a transposition event of the Ty element *YERCTy1-1* from chromosome V to chromosome III, adjacent to the previously annotated *YCLWTy2-1* located 7.3 kb to the left side of the HO recognition site. Long-terminal repeat (LTR)-retrotransposons are widely spread in eukaryotic genomes and are thought to be the evolutionary progenitors of retroviruses. These retrotransposons elements are located in extensively heterochromatic silenced genomic regions because, although their transposition has been suggested to, in some cases, enhance genomic plasticity or regulate the expression of adjacent genes, potentially benefiting cellular fitness, their uncontrolled transposition could be highly detrimental for genome integrity. In this scenario, it is plausible that the presence of heterochromatic sequences nearby the HO cleavage site might hinder end-resection in cells lacking PP4^Psy2^ activity. Supporting this hypothesis, disruption of both *YERCTy1-1* and *YCLWTy2-1* elements improves resection of *psy2Δ* cells. This effect is particularly significant in the context of tandem Ty elements, since in a strain harboring a unique recombinant N-*YERCTy1-1/*C-*YCLWTy2-1* Ty element adjacent to the HO site, the influence of PP4^Psy2^ was minimal. However, a slight delay in resection was still detectable, likely attributable to the intrinsic challenges associated with resecting repetitive genomic regions (Campos *et al*., 2023; Ramos *et al*., 2022). This retrotransposon-dependent negative effect extends beyond DNA end resection to impact other cellular processes. It has been reported that Ty elements repress homologous recombination at meiotic hotspots through the establishment of a closed chromatin structure, which relies on the N-terminal sequences flanking the Ty elements (Ben-Aroya *et al*., 2004). Moreover, two adjacent inverted Ty elements can form long hairpin structures on the lagging strand during replication, leading to a region of genetic instability (Casper *et al*., 2009). Alternatively, retrotransposons elements tend to accumulate R-loops (Zeng *et al*., 2021), which are associated with genomic instability and replication stress, likely due to an impeded DNA polymerase progression and replication fork stalling (Aguilera & Garcia-Muse, 2012). Whether retrotransposons halt resection progression through the accumulation of R-loops is an intrigued question for future research. In this context, the finding that RNaseH overexpression leads to excessive strand resection (Ohle *et al*, 2016) suggests a potential mechanism contributing to this phenomenon.

How does PP4^Psy2^ enhance resection through retrotransposons elements? It is likely that the elevated levels of Rad53 inherent of *psy2Δ* cells might lead to misregulation of downstream targets involved in the processing of the DNA lesion. Helicases are ideal candidates for facilitating resection through genomic heterochromatic regions. The Rrm3 helicase unwinds DNA with a 5’-3’ polarity and was originally identified as a suppressor of mitotic recombination in tandem DNA arrays (Keil & McWilliams, 1993). Rrm3 assists replication fork progression through barriers at rDNA loci (Ivessa *et al*, 2000) and tRNA genes (Osmundson *et al*, 2017). In the DDR context, Rrm3 has been implicated in repairing replication-induced DNA breaks via sister chromatid recombination (Munoz-Galvan *et al*, 2017) and in DNA repair in cells lacking Srs2 or Sgs1 helicases, likely due to the complementary polarities of their helicase activities (Schmidt & Kolodner, 2004). Additionally, Rrm3 has been associated to Ty1 transposition, possibly by regulating certain stages of the HR pathway (Scholes *et al*, 2001). Another key helicase involved in DNA repair is Pif1, a broadly conserved 5’-3’ helicase essential for the maintenance of genome integrity. Pif1 plays diverse roles, including telomere length regulation (Schulz & Zakian, 1994; Zhou *et al*, 2000), Okazaki fragment maturation (Budd *et al*, 2006; Rossi *et al*, 2008), break-induced replication (Li *et al*, 2021; Vasianovich *et al*, 2014), and aiding replication fork progression and resection through genomic regions sites containing G-quadruplexes (Duan *et al*, 2015; Jimeno *et al*, 2018; Paeschke *et al*, 2011). Importantly, Mec1-Rad53-dependent phosphorylation of Pif1 inhibits telomerase activity at DSBs (Makovets & Blackburn, 2009) and direct it to telomeres (Vasianovich *et al*., 2014). Furthermore, Rad53-mediated phosphorylation of both Pif1 and Rrm3 inhibits their helicase activity to prevent aberrant fork transitions under replicative stress (Rossi *et al*., 2015). In this context and given the high levels of Rad53 phosphorylation observed in *psy2Δ* cells in response to an HO break, it is plausible that the activities of these helicases are downregulated in this mutant, contributing to the resection delay at the *YERCTy1-1/YCLWTy2-1* region and the asymmetric resection profile observed. Supporting this notion, we found that Rrm3 and Pif1 are persistently phosphorylated in the absence of PP4^Psy2^ activity during the DSB response. The role of Rrm3 and Pif1 in SSA seems to be redundant, as the individual depletion of either helicase causes only a minor defect in resection compared to the pronounced defect observed in the double *pif1Δ rrm3Δ* mutant. This redundancy suggests that both helicases cooperate to facilitate resection through Ty elements. Consistent with this, the absence of either helicase in *psy2Δ* cells does not result in additive defects in resection progression, aligning with the hypothesis that PP4^Psy2^ regulates both helicases in response to DNA damage. Importantly, overexpression of Rrm3, but not a nuclear-localized version of Pif1, rescues the resection defects of *psy2Δ* cells in SSA. This result confirms that impaired resection through heterochromatic Ty regions in *psy2Δ* cells is due to defective Rrm3 function, and hints that excessive levels of Pif1 might be detrimental for resection progression. Interestingly, this effect is only observed distantly from the HO break, as the HO cleavage product is timely processed under Pif1-m1 overexpression. This suggests that high Pif1 levels are incompatible with extended resection while efficiently sustaining short-range processing. Nevertheless, this effect occurs independently of the presence of retrotransposons elements nearby the break sites, as is observed on both sides of the HO break.

In summary, we propose that persistent Rad53 phosphorylation in the absence of PP4^Psy2^ activity leads to elevated H2A phosphorylation, which in turn increases the accumulation of chromatin-bound Rad9. This cascade negatively impacts DNA resection, with the effect being more pronounced in heterochromatic regions containing joint retrotransposon sequences. Efficient resection through these regions requires fully active Pif1 and Rrm3 helicases to facilitate the processing of the DNA break to enable SSA. It is important to remark that, given the vast number of functions attributed to Rrm3 and Pif1, it is possible that their PP4-dependent regulation might be extended to other physiological processes. Indeed, PP4 and Rrm3/Pif1 share several overlapping molecular functions, including the resolution of joint molecules (Jablonowski *et al*, 2015), stabilization of replication forks (O’Neill *et al*., 2004; Rossi *et al*., 2015) and regulation of telomerase activity at DSBs (Makovets & Blackburn, 2009; Vasianovich *et al*., 2014; Zhang & Durocher, 2010). This overlap raises the interesting possibility that PP4 might exert a broader control over Pif1 and Rrm3 in various cellular contexts. Further exploration of these interactions might uncover novel mechanisms by which PP4 modulates the activity, stability, or localization of these helicases, thereby contributing to genome maintenance. Additionally, such studies could reveal broader implications of PP4 in cellular processes beyond its known functions, offering deeper insights into its biological significance.

## Structured Methods

### Yeast strains and growing conditions

The strains utilized in this study are detailed in Supplementary Table 1. Strain YMV80 (Ramos *et al*., 2022), generously provided by J. Haber (Brandeis University, Waltham, MA, USA) was included. Gene disruption and tagging were performed by gene targeting using PCR products, as previously described (Goldstein & McCusker, 1999; Janke *et al*, 2004), with oligonucleotides listed in Supplementary Table 3. Strains derived from YMV80 were induced by adding 2% galactose to cells cultured in YP with 2% raffinose. Samples for DNA analysis were collected both before and after the addition of galactose to the media. G1 phase arrest was achieved by using the alpha factor pheromone at a concentration of 3 μM. All strains containing the *hta1/hta2-S129\** versions of H2A were constructed as previously described (Downs *et al*., 2000). Strains and plasmids used in this study are available upon request through the lead contact.

### Chemiluminescent Southern blotting

10 ml of cell cultures at an OD_600_ of 0.4 were harvested by centrifugation and washed with 1 ml of PBS. The pellets were then flash-frozen and stored at -80°C. For cell lysis, the pellets were treated with 40 units of lyticase in DNA preparation buffer (1% SDS, 100 mM NaCl, 50 mM Tris-HCl, 10 mM EDTA) for 10 min. DNA extraction was performed by incubating with phenol:chloroform:isoamylalcohol (25:24:1) for 10 min. After centrifugation, the aqueous phase was precipitated with ethanol and resuspended in TE buffer. Genomic DNA was digested with *Sty*I or *Kpn*I, separated on 1% agarose gels, and subjected to Southern blotting. Probes were generated by labeling a PCR-amplified DNA fragment with a nucleotide mix containing fluorescein-12-dUTP (Fisher Scientific, 10354280). The oligonucleotides used for probes synthesis are listed in Supplementary Table 2. Detection was carried out using an anti-fluoresceine antibody conjugated to alkaline phosphatase (F(a,b)2 fluorescein polyclonal antibody AP, Life Technology, 700-105-096) at a 1:200,000 dilution. Membranes were incubated with CDP-Star detection reagent (Cytiva, RPN3682) and exposed to films. Imaging processing and analysis were performed using the FIJI software (https://imagej.net/Fiji).

### Western blots

Samples for western blotting were prepared using trichloroacetic acid (TCA) extraction. Cells were collected, centrifugated, and fixed with 20% TCA. Cell lysates were obtained by breaking the cells with glass beads in a FastPrep homogenizer (MPBio) for three 20-second cycles at a power setting of 5.5. Proteins were precipitated by centrifuging at 2,200 g for 5 minutes at 4°C, and the pellet was solubilized in 1 M Tris-HCl (pH8) and SDS-PAGE loading buffer. After boiling for 10 min, the insoluble material was removed by centrifugation, and the supernatant was loaded onto 6% acrylamide gels. To facilitate the separation of Psy2, Pif1 and Rrm3 phosphobands, a final concentration of 10 μM, 20 μM and 10 μM of Phos-Tag was used, respectively (Wako, 300-93523). Proteins were transferred to a PVDF membrane (Hybond-P, GE Healthcare, 15259894) and blocked with 5% milk in PBS-Tween (0.1%). Anti-Rad53 antibody (Abcam, ab104232) and anti-Histone H2A (phospho-S129) antibody (Abcam, ab15083) were used at a 1:2,000 dilution and 1:500 respectively. The secondary anti-rabbit antibody was used at a 1:5,000 dilution (GE Healthcare, NA934). Anti-MYC antibody for Psy2-9MYC (Merk, C3956) was used at a 1:2500. The secondary anti-rabbit antibody was used at a 1:5,000 dilution (GE Healthcare, NA934). Anti-HA antibody for Pif1-HA (Anti-HA 12CA5, Merk, 11583816001) was used at a 1:2,500 dilution. Anti-HA antibody for Rrm3-HA (Anti-HA High Affinity, Merk, 11867423001) was used at a 1:2,000 dilution. The secondary anti-mouse and anti-rat antibody were used at a concentration of 1:25,000 and 1:10,000, respectively (GE Healthcare, NA931 and NA935). After washing the membranes with PBS-Tween, they were incubated with SuperSignal^®^ West Femto (Thermo Scientific, 10391544) and exposed to Amersham^TM^ Hyperfilm^TM^ ECL (Cytiva, 28-9068-37).

### Genomic sequencing and data analysis

#### Short-read sequencing and data analysis

For short-reads genomic analyses of *rad51Δ* and *rad51Δ psy2Δ* cells in the *YMV80_v2* background, 1μgr genomic DNA was used for library preparation and subsequently sequenced using a DNBSEQ G400 platform in PE150 (BGI). Briefly, genomic DNA was processed through fragmentation, end-repair, and 3’-adenylation reactions. Subsequently, adaptors were ligated to the 3’-adenylated fragments. PCR was performed to amplify the adaptor-ligated products. Following this, a circularization reaction was configured, resulting in the formation of single-stranded cyclized DNA molecules, that were amplified using rolling circle amplification producing DNA nanoballs (DNBs). Sequencing was performed using the combinatorial probe-anchor synthesis (cPAS) method. Reads containing adapters sequences or low-quality bases were excluded from the analysis. This processing was conducted using SOAPnuke, a tool developed by BGI, with the following parameters: -n 0.001 -l 20 -q 0.4 --adaMis 3 --rmdup –minReadLen 150.

For DNA samples of *rad51Δ psy2Δ* cells in the *YMV80_v3* background, genomic libraries were generated by using the Nextera XT DNA Library Preparation Kit (Illumina). Briefly, 10 ngr of DNA from the genomic extraction (described above) was incubated at 55°C for 5 minutes in an enzymatic tagmentation reaction to fragment the DNA and add adapter sequences. The tagmented DNA was then amplified using a 7-cycle PCR program to incorporate indexes and sequences necessary for cluster formation. DNA libraries were purified with Sera-Mag® magnetic beads (Cytiva, 29343052) and assessed on an Agilent 2100 Bioanalyzer using a High Sensitivity chip. Library concentrations were measured using a Qubit 3.0 fluorometer. A genomic DNA pool (2.1pM) comprising all libraries, along with 1.8% PhiX library as a control, was sequenced on an Illumina NextSeq500 platform using a paired-end 75 bp protocol.

Sequenced reads were aligned to the *S. cerevisiae YMV80_v2* or *YMV80_v3* reference genomes (Supplementary Data 1 and 2, YMV80_v2_RG.fasta and YMV80_v3_RG.fasta) using Bowtie (v1.0.0) (Langmead *et al*, 2009) with the -v 0 alignment mode (allowing no mismatches), and the -m 1 reporting mode (suppressing alignments if more than one reportable alignments was found for a read). The resulted SAM files were converted to BAM files using Samtools (v1.12) (Li *et al*, 2009). BAM data were normalized using Reads Per Genomic Content (RPGC) with the Deeptools utility (v3.5.1). BedGraph files generated from these alignments were used to calculate chromosomal coverage in the *YMV80* background strains with Bedtools (v2.30.0). Coverage profiles were visualized using the Integrative Genome Browser IGB (v9.0.2) (Freese *et al*, 2016) and IGV (v2.16.2) (Robinson *et al*, 2011).

#### Long-read sequencing and analysis

For long-read *de novo* genome assembly, libraries were prepared using 200 fmol of genomic DNA using the sequencing kit SQK-NBD114-24 (Oxford Nanopore Technologies) in accordance with the manufacturer’s protocol. The prepared libraries were loaded onto a FLO-MIN114 flow cell and sequenced using a MinION Mk1C device (Oxford Nanopore Technologies) with MinKNOW software (v23.07.12), following the manufacturer’s guidelines.

Long-reads sequences were subjected to quality filtering and trimming using the Filtlong tool (v0.2.1), which selectively retains high-quality reads based on length (--min_length 1000) and quality score (--min_mean_q 10) thresholds to improve downstream analysis. The filtered reads were then assembled *de novo* using Flye assembler (v2.9.5), which is specifically designed for long-read data. The resulting contigs were further refined and corrected using Racon (v1.5.0), a polishing tool that employs alignment data to enhance the accuracy of consensus sequences by correcting errors associated with insertions, deletions and substitutions.

#### Generation of Read Coverage Profiles

To normalize 2D/3D read coverage profiles and colormaps, we adopted the approach described in (Campos *et al*., 2023). Briefly, the coverage was calculated by averaging reads counts over 0.3 kb sections along the chromosomes. Coverage values of different time point samples after HO induction were normalized against the corresponding sections from the 0 h sample (T0). Additionally, normalization was performed relative to a 120 kb region on chromosome V (coordinates 210000-330000) using the formula *C_N_ = [(CIII_Tx_ / CIII_T0_) / (CV_Tx_ / CV_T0_)]*, where *C_N_* is the normalized coverage, *CIII_Tx_* and *CIII_T0_* represent the averaged coverage of each 0.3 kb sections from chromosome III at time points T_x_ and T_0_, respectively, and *CV_Tx_* and *CV_T0_* denote the averaged coverage of the 120 kb section on chromosome V at the same respective time points. Colormaps were generated by translating normalized read coverage values from the 3D profiles into a gradient of colors, ranging from red (indicating high coverage) to purple (indicating low coverage).

#### Analysis of DNA end resection and re-synthesis dynamics

To quantitatively measure resection rates, we first averaged the coverage of 300 nt fractions across a +/-30 kb region surrounding the HO site for each time-point assessed in the *rad51Δ _v2* and *rad51Δ psy2Δ_v2*. This value was reduced to -22 kb at the left side of the HO break in the *rad51Δ psy2Δ_v3* to -22 kb due to the shorter inter-*U2* distance. Using the trend line equations generated for each fraction, we determined the time required for coverage to decrease from 1 to 0.5. This measurement indicates the average time that a cell population needs to reach a particular distance from the HO break. In SSA, the DNA region flanked by the *U2* homologous sequences does not undergo DNA re-synthesis. Therefore, the decrease in coverage in this region exclusively reflects the progression of resection, allowing us to use a 0-15 h time interval to generate the coverage trendlines equations when dissecting the left side of the HO break. For the right side of the HO site, where coverage is influenced by both resection and DNA re-synthesis, we limited the analysis to the 0-6 h interval, thus minimizing the impact of DNA re-polymerization on the assessment of resection progression. The resulting values for each 300 nt fraction were plotted as graphs representing time (h) versus distance (kb). To calculate the resection rates, the left and right sides of the graph were analyzed separately. Using the trend line equations we determine the time (*y-*value) required for resection to cover specific distances (*x-*values) from the HO break. Resection rate was then calculated using the formula *v = Δs/Δt*, where *Δs* represents the distance covered and *Δt* the corresponding time.

Synthesis rate was calculated by using a strategy similar to that employed for resection. We analyzed the averaged coverages of 300 nt fractions spanning from the HO site to 10 kb to the right. Using the trend line equations generated for each fraction, we determined the time required for coverage to recover from resection and reach a value of 0.5 within the 9-24 h interval. This value represents the average time for the cell population to restore the resected DNA at a specific position relative to the HO break. The resulting values for each 300 nt fraction were then plotted as graphs of time (h) *versus* distance (kb).

#### Statistics and reproducibility

All Southern blots and genome-wide sequencing experiments in this study were conducted with at least two biological replicates to ensure the result reproducibility. Statistical analyses were performed using GraphPad Prism software, with *P*-values calculated using a two-tailed unpaired Student′s *t-*test. A significance threshold of *P* ≤ 0.05 was applied across all experiments. Detailed information on the number of replicates, statistical tests used, and *P-*values for each analysis is provided in the corresponding figure legends.

## Acknowledgments

We thank P. San Segundo, J. Haber and J.A. Downs for providing valuable strains and plasmids. We particularly thank P. San Segundo, C. Martín-Castellanos, J. Matos and F. Machín for insightful discussions and valuable feedback on the work. We also extend our gratitude to the members of our laboratories for their constructive discussions and support. This work was funded by projects BFU2016-77081-P, PGC2018-097963-B-100, and PID2021-125290NB-I00, granted by the MCIN/AEI/10.13039/501100011033/ and co-financed by “FEDER, Una manera de hacer Europa”, awarded to A. C-B. Partial institutional support for the IBFG was provided by the “Junta y Castilla y León” through the “Escalera de Excelencia” program (Ref. CLU-2017-03), co-funded by P.O. FEDER de Castilla y León 2014-2020. A.C. received a predoctoral fellowship from the “Junta de Castilla y León”. L.I. was supported by a “Jae-INTRO” grant from the CSIC and is currently recipient of predoctoral fellowship from the “Junta de Castilla y León”. C.D. was funded by a predoctoral fellowships from the “Universidad de Salamanca” and is currently beneficiary of a FPU from the Spanish Ministry of Education.

## Author Contributions

A.C, L.P., L.I., C.D. M.V and A. C-B. contributed to performing all the experiments. L.I. assisted with the analysis of the sequencing data. M.S. participated to the creation of the YMV80_v2 and YMV80_v3 reference genomes. A.C-B. designed the research plan, supervised the experiments, collaborated in the analysis of the data, and wrote the manuscript. All authors discussed the results and approved the final version of the manuscript.

## Disclosure and competing interest statement

The authors declare no competing interests.

## Data Availability Section

### Data Availability

Genomic datasets produced in this study are available in the Sequence Read Archive (SRA) database at the link: https://dataview.ncbi.nlm.nih.gov/object/PRJNA1208738?reviewer=ji4pdr9n343tull82n938nc5p9

This paper does not report original code. Further information and requests required to reanalyze the data reported in this study is available from the lead contact upon request, Andrés Clemente Blanco.

## Tables

**Table 1.**
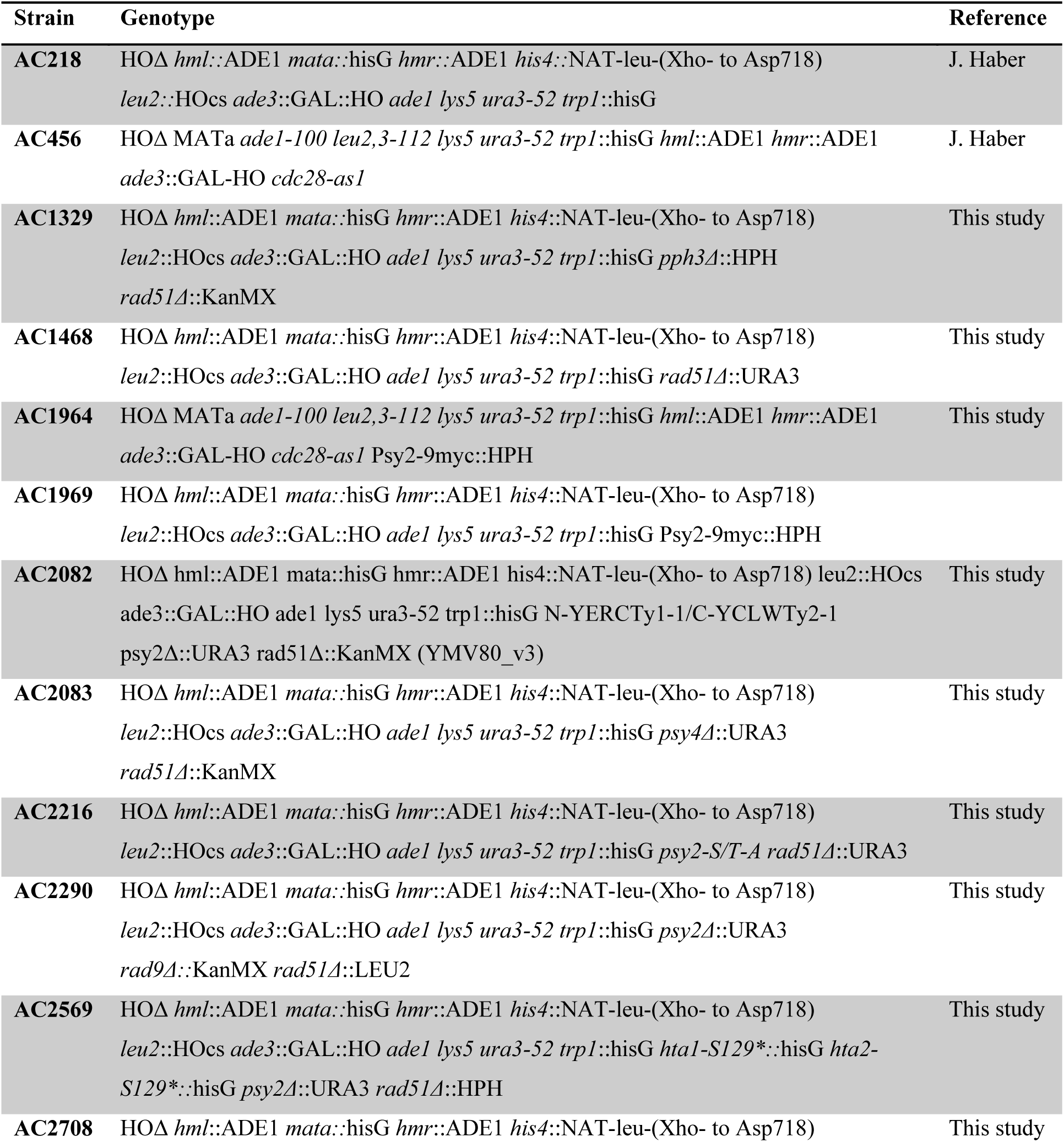
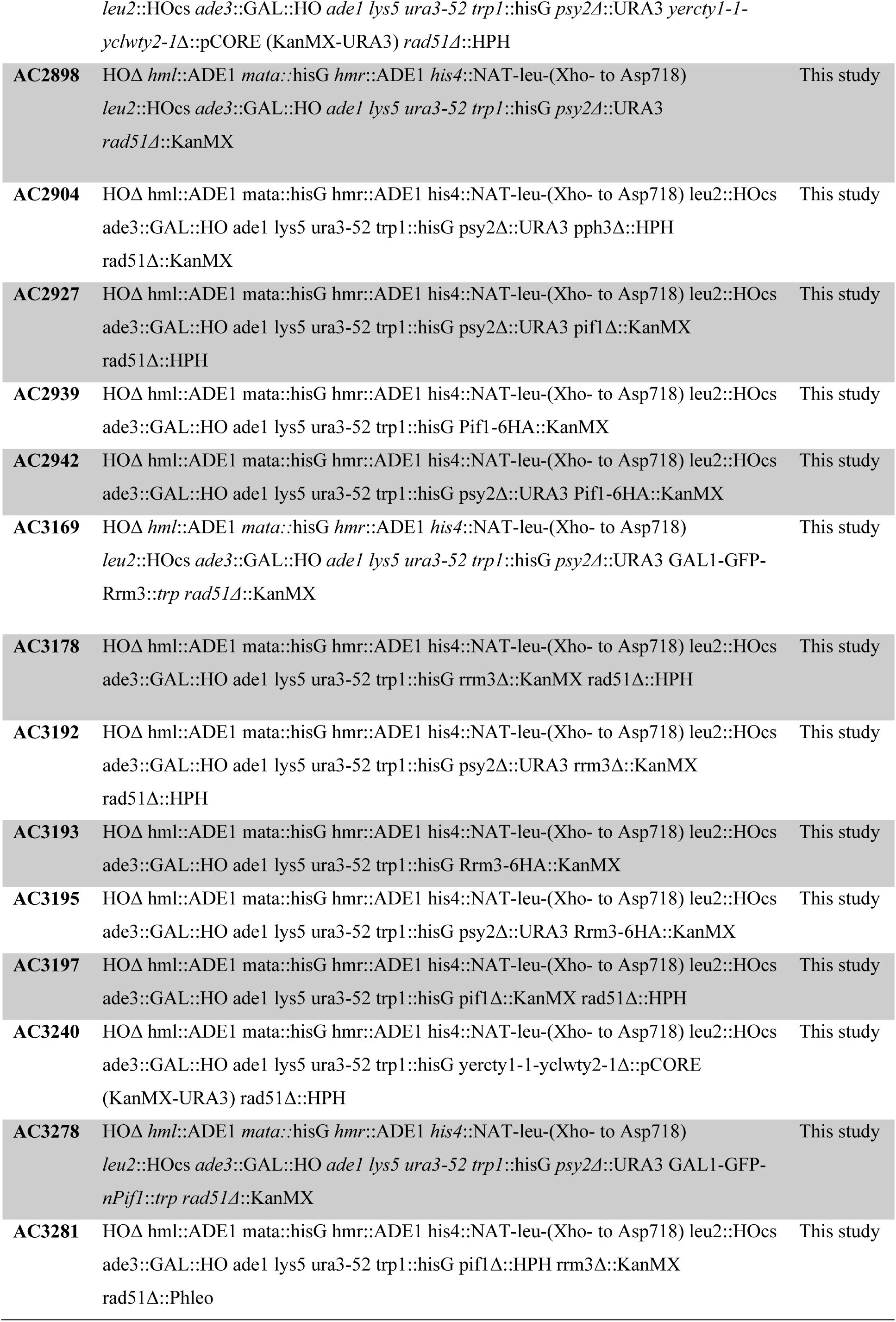
Genotypes of strains used in this study.

| Strain | Genotype | Reference |
| --- | --- | --- |
| **AC218** | HOΔ *hml::ADE1 mata::hisG hmr::ADE1 his4::NAT-leu*-(Xho- to Asp718)<br>*leu2::HOcs ade3::GAL::HO ade1 lys5 ura3-52 trp1::hisG* | J. Haber |
| **AC456** | HOΔ MATa *ade1-100 leu2,3-112 lys5 ura3-52 trp1::hisG hml::ADE1 hmr::ADE1*<br>*ade3::GAL-HO cdc28-as1* | J. Haber |
| **AC1329** | HOΔ *hml::ADE1 mata::hisG hmr::ADE1 his4::NAT-leu*-(Xho- to Asp718)<br>*leu2::HOcs ade3::GAL::HO ade1 lys5 ura3-52 trp1::hisG pph3Δ::HPH*<br>*rad51Δ::KanMX* | This study |
| **AC1468** | HOΔ *hml::ADE1 mata::hisG hmr::ADE1 his4::NAT-leu*-(Xho- to Asp718)<br>*leu2::HOcs ade3::GAL::HO ade1 lys5 ura3-52 trp1::hisG rad51Δ::URA3* | This study |
| **AC1964** | HOΔ MATa *ade1-100 leu2,3-112 lys5 ura3-52 trp1::hisG hml::ADE1 hmr::ADE1*<br>*ade3::GAL-HO cdc28-as1 Psy2-9myc::HPH* | This study |
| **AC1969** | HOΔ *hml::ADE1 mata::hisG hmr::ADE1 his4::NAT-leu*-(Xho- to Asp718)<br>*leu2::HOcs ade3::GAL::HO ade1 lys5 ura3-52 trp1::hisG Psy2-9myc::HPH* |  |
| **AC2082** | HOΔ *hml::ADE1 mata::hisG hmr::ADE1 his4::NAT-leu*-(Xho- to Asp718) *leu2::HOcs*<br>*ade3::GAL::HO ade1 lys5 ura3-52 trp1::hisG N-YERCTy1-1/C-YCLWTy2-1*<br>*psy2Δ::URA3 rad51Δ::KanMX (YMV80_v3)* | This study |
| **AC2083** | HOΔ *hml::ADE1 mata::hisG hmr::ADE1 his4::NAT-leu*-(Xho- to Asp718)<br>*leu2::HOcs ade3::GAL::HO ade1 lys5 ura3-52 trp1::hisG psy4Δ::URA3*<br>*rad51Δ::KanMX* | This study |
| **AC2216** | HOΔ *hml::ADE1 mata::hisG hmr::ADE1 his4::NAT-leu*-(Xho- to Asp718)<br>*leu2::HOcs ade3::GAL::HO ade1 lys5 ura3-52 trp1::hisG psy2-S/T-A rad51Δ::URA3* | This study |
| **AC2290** | HOΔ *hml::ADE1 mata::hisG hmr::ADE1 his4::NAT-leu*-(Xho- to Asp718)<br>*leu2::HOcs ade3::GAL::HO ade1 lys5 ura3-52 trp1::hisG psy2Δ::URA3*<br>*rad9Δ::KanMX rad51Δ::LEU2* | This study |
| **AC2569** | HOΔ *hml::ADE1 mata::hisG hmr::ADE1 his4::NAT-leu*-(Xho- to Asp718)<br>*leu2::HOcs ade3::GAL::HO ade1 lys5 ura3-52 trp1::hisG hta1-S129\*::hisG hta2-*<br>*S129\*::hisG psy2Δ::URA3 rad51Δ::HPH* | This study |
| **AC2708** | HOΔ *hml::ADE1 mata::hisG hmr::ADE1 his4::NAT-leu*-(Xho- to Asp718) | This study |
| *leu2::HOcs ade3::GAL::HO ade1 lys5 ura3-52 trp1::hisG psy2Δ::URA3 yercty1-1-yclwty2-1Δ::pCORE (KanMX-URA3) rad51Δ::HPH* |  |  |
| **AC2898** | *HOΔ hml::ADE1 mata::hisG hmr::ADE1 his4::NAT-leu-(Xho- to Asp718) leu2::HOcs ade3::GAL::HO ade1 lys5 ura3-52 trp1::hisG psy2Δ::URA3 rad51Δ::KanMX* | This study |
| **AC2904** | *HOΔ hml::ADE1 mata::hisG hmr::ADE1 his4::NAT-leu-(Xho- to Asp718) leu2::HOcs ade3::GAL::HO ade1 lys5 ura3-52 trp1::hisG psy2Δ::URA3 pph3Δ::HPH rad51Δ::KanMX* | This study |
| **AC2927** | *HOΔ hml::ADE1 mata::hisG hmr::ADE1 his4::NAT-leu-(Xho- to Asp718) leu2::HOcs ade3::GAL::HO ade1 lys5 ura3-52 trp1::hisG psy2Δ::URA3 pif1Δ::KanMX rad51Δ::HPH* | This study |
| **AC2939** | *HOΔ hml::ADE1 mata::hisG hmr::ADE1 his4::NAT-leu-(Xho- to Asp718) leu2::HOcs ade3::GAL::HO ade1 lys5 ura3-52 trp1::hisG Pif1-6HA::KanMX* | This study |
| **AC2942** | *HOΔ hml::ADE1 mata::hisG hmr::ADE1 his4::NAT-leu-(Xho- to Asp718) leu2::HOcs ade3::GAL::HO ade1 lys5 ura3-52 trp1::hisG psy2Δ::URA3 Pif1-6HA::KanMX* | This study |
| **AC3169** | *HOΔ hml::ADE1 mata::hisG hmr::ADE1 his4::NAT-leu-(Xho- to Asp718) leu2::HOcs ade3::GAL::HO ade1 lys5 ura3-52 trp1::hisG psy2Δ::URA3 GAL1-GFP-Rrm3::trp rad51Δ::KanMX* | This study |
| **AC3178** | *HOΔ hml::ADE1 mata::hisG hmr::ADE1 his4::NAT-leu-(Xho- to Asp718) leu2::HOcs ade3::GAL::HO ade1 lys5 ura3-52 trp1::hisG rrm3Δ::KanMX rad51Δ::HPH* | This study |
| **AC3192** | *HOΔ hml::ADE1 mata::hisG hmr::ADE1 his4::NAT-leu-(Xho- to Asp718) leu2::HOcs ade3::GAL::HO ade1 lys5 ura3-52 trp1::hisG psy2Δ::URA3 rrm3Δ::KanMX rad51Δ::HPH* | This study |
| **AC3193** | *HOΔ hml::ADE1 mata::hisG hmr::ADE1 his4::NAT-leu-(Xho- to Asp718) leu2::HOcs ade3::GAL::HO ade1 lys5 ura3-52 trp1::hisG Rrm3-6HA::KanMX* | This study |
| **AC3195** | *HOΔ hml::ADE1 mata::hisG hmr::ADE1 his4::NAT-leu-(Xho- to Asp718) leu2::HOcs ade3::GAL::HO ade1 lys5 ura3-52 trp1::hisG psy2Δ::URA3 Rrm3-6HA::KanMX* | This study |
| **AC3197** | *HOΔ hml::ADE1 mata::hisG hmr::ADE1 his4::NAT-leu-(Xho- to Asp718) leu2::HOcs ade3::GAL::HO ade1 lys5 ura3-52 trp1::hisG pif1Δ::KanMX rad51Δ::HPH* | This study |
| **AC3240** | *HOΔ hml::ADE1 mata::hisG hmr::ADE1 his4::NAT-leu-(Xho- to Asp718) leu2::HOcs ade3::GAL::HO ade1 lys5 ura3-52 trp1::hisG yercty1-1-yclwty2-1Δ::pCORE (KanMX-URA3) rad51Δ::HPH* | This study |
| **AC3278** | *HOΔ hml::ADE1 mata::hisG hmr::ADE1 his4::NAT-leu-(Xho- to Asp718) leu2::HOcs ade3::GAL::HO ade1 lys5 ura3-52 trp1::hisG psy2Δ::URA3 GAL1-GFP-nPif1::trp rad51Δ::KanMX* | This study |
| **AC3281** | *HOΔ hml::ADE1 mata::hisG hmr::ADE1 his4::NAT-leu-(Xho- to Asp718) leu2::HOcs ade3::GAL::HO ade1 lys5 ura3-52 trp1::hisG pif1Δ::HPH rrm3Δ::KanMX rad51Δ::Phleo* | This study |

**Table 2.**
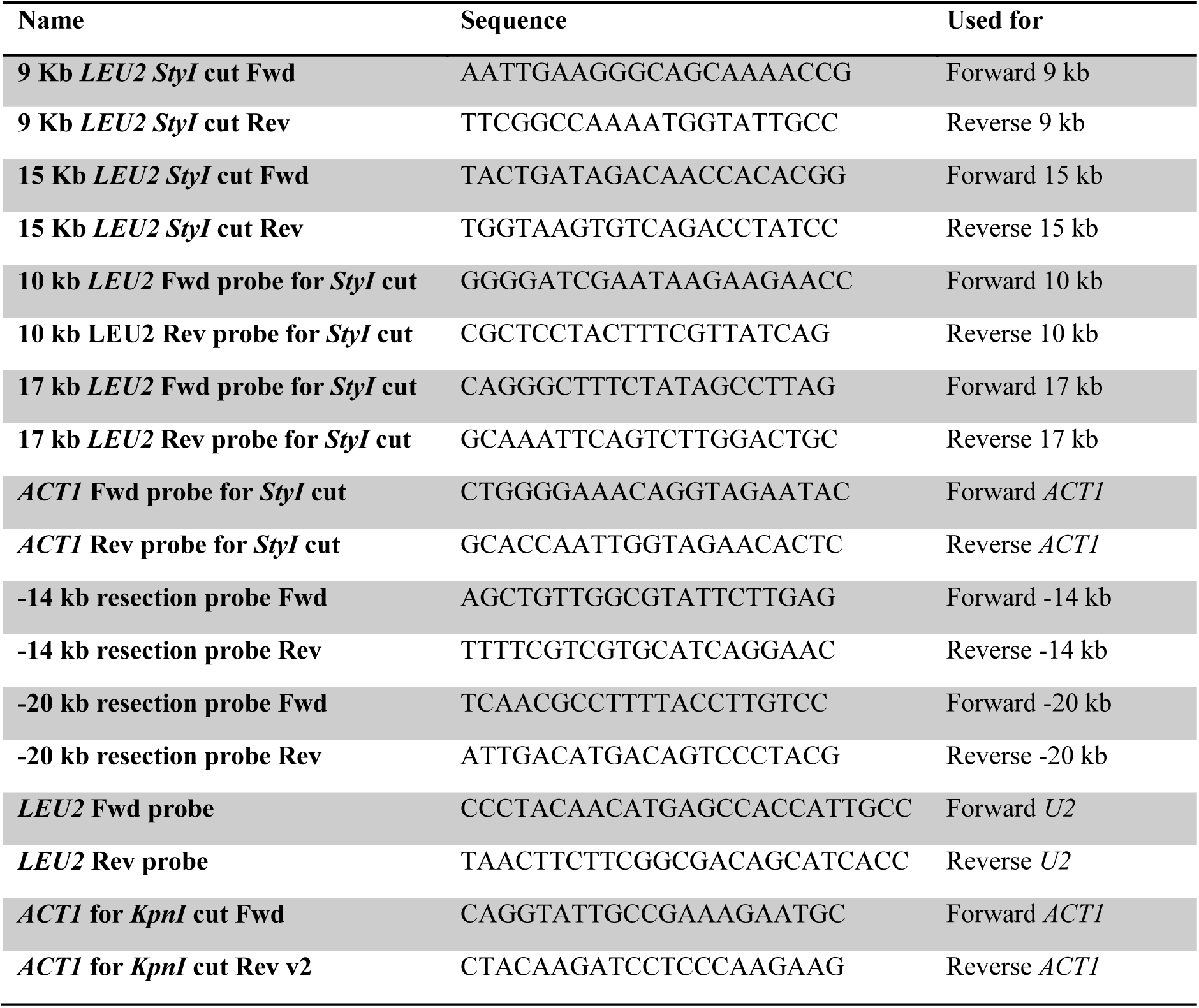
Oligonucleotides used for probes synthesis.

**Table 3.**
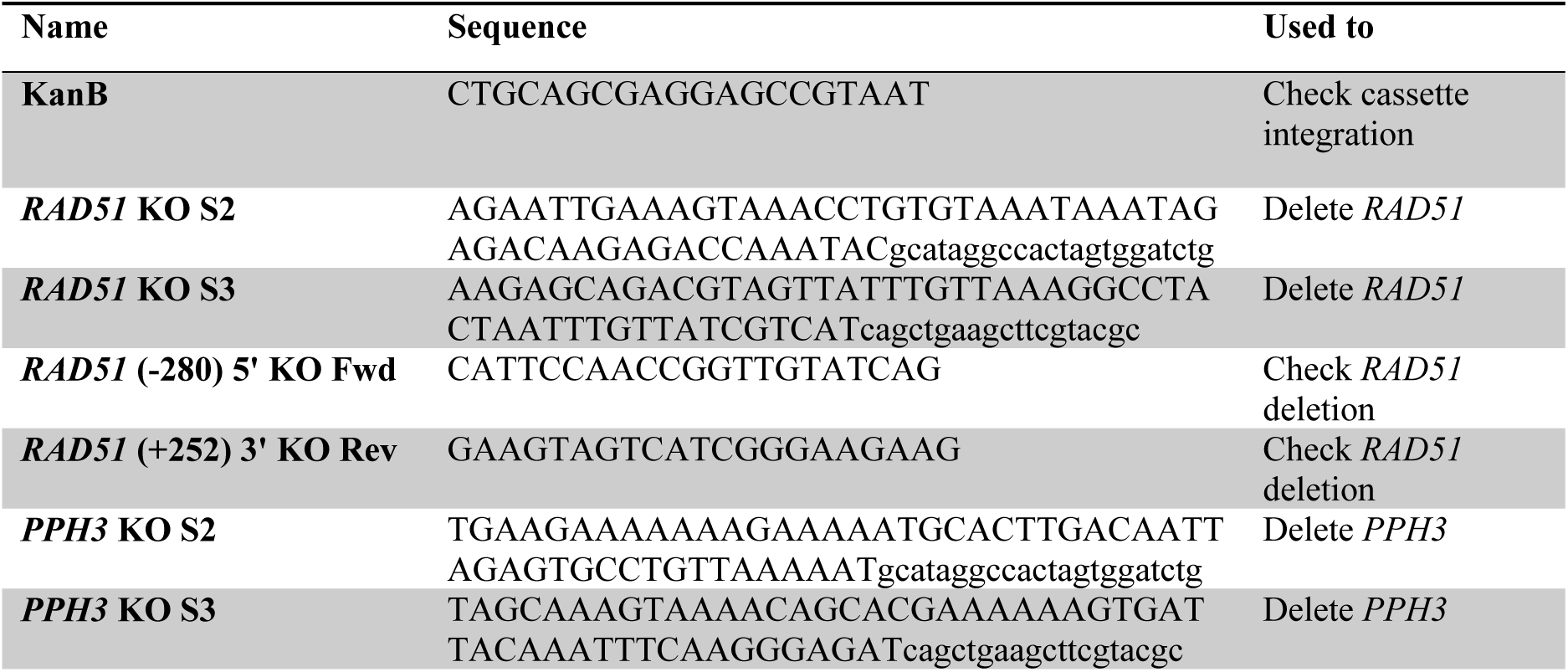
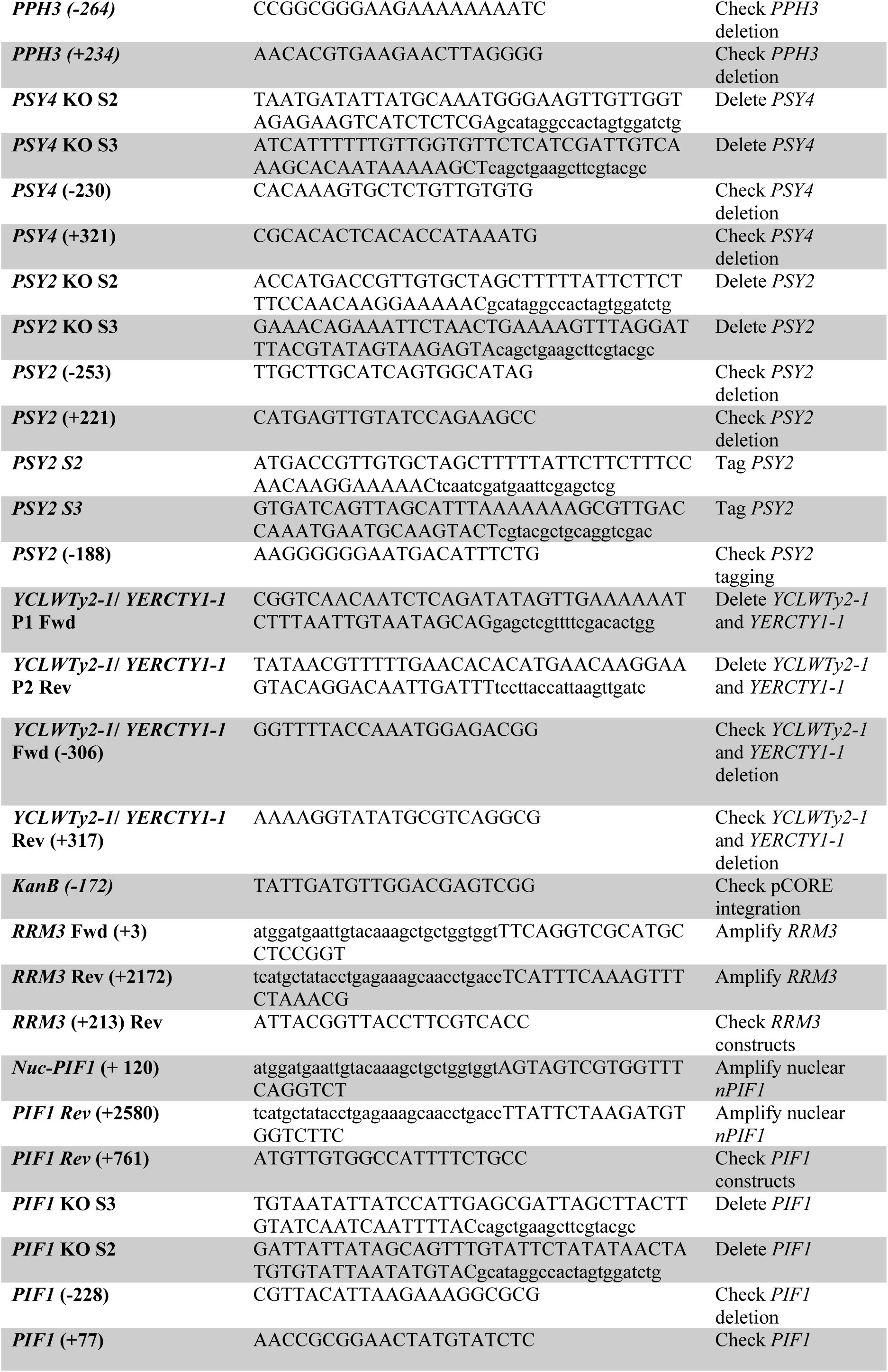
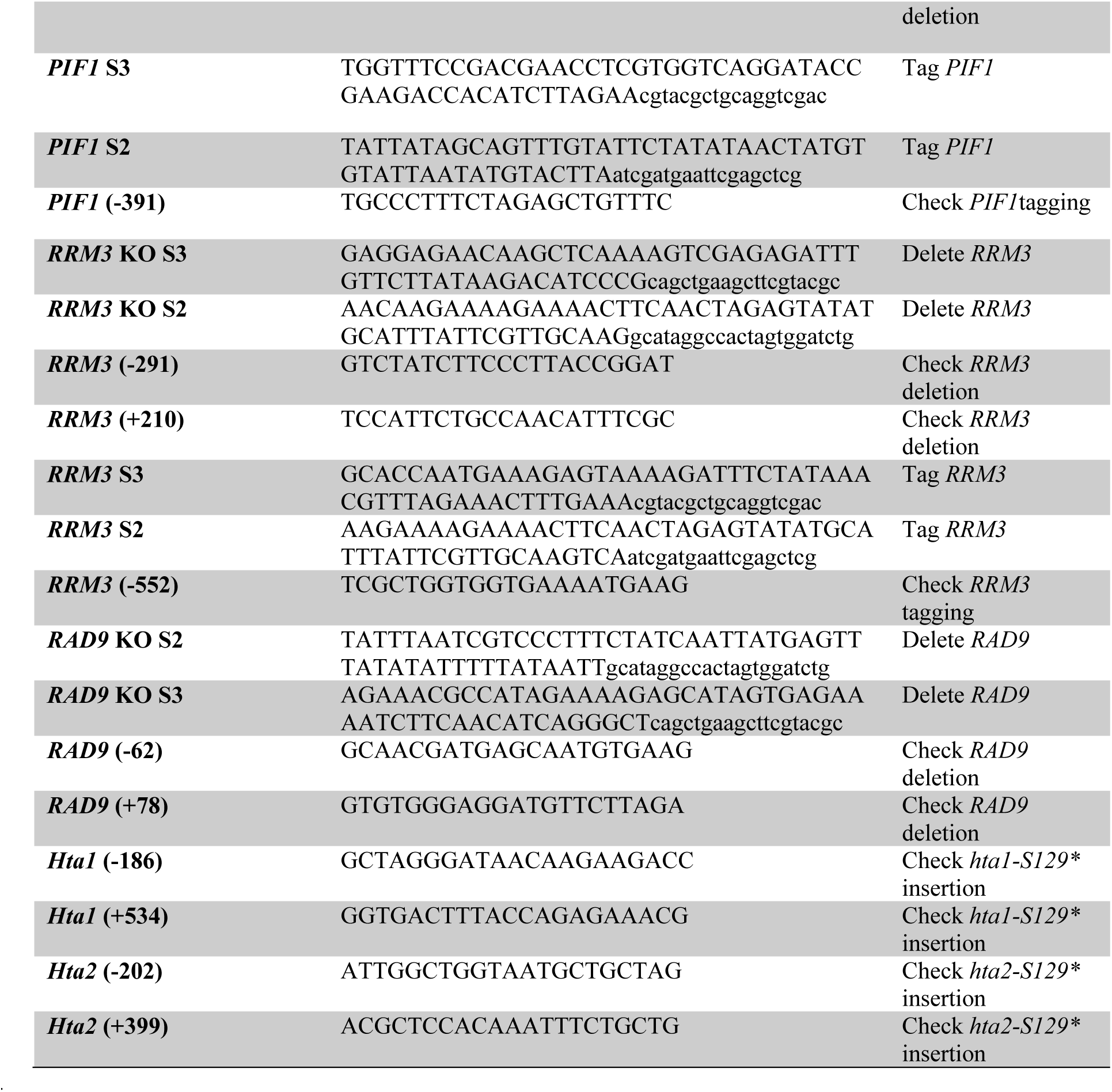
Other oligonucleotides used in this study.

## Expanded View Figure Legends

**Expanded View Figure 1.**
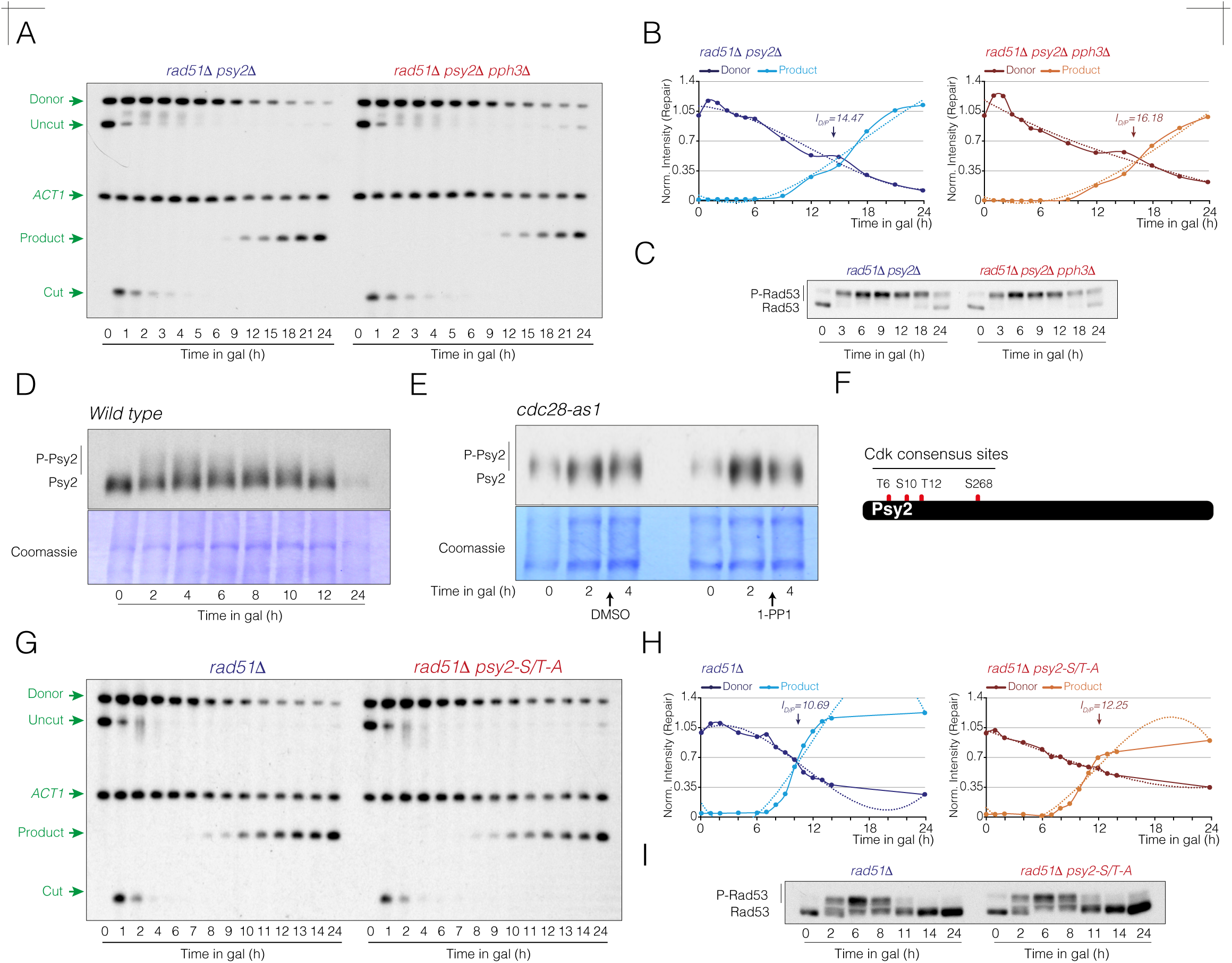
Psy2 and Pph3 display redundant roles in SSA repair. **A.** Southern blot analysis of *rad51Δ psy2Δ* and *rad51Δ psy2Δ pph3Δ* strains. Cells were grown overnight in YP-Raffinose before expressing the HO nuclease by adding galactose to the medium. Samples were collected at the indicated time points, genomic DNA was extracted, digested with *Kpn*I and analyzed by Southern blot using the probes shown in figure 1A. The *ACT1* gene sequence was used as loading control. **B.** Graphs show the quantification of the donor and product band signals from A, averaged from two biological replicates and normalized to their respective *ACT1* and uncut T0 signals. The intersection point of the donor and product trend lines (I_D/P_ value) represents the mean from two independent experiments. The statistical significance (*P*-value) of differences between both strains assessed by a two-tailed unpaired Student’s *t-*test is 0.0153 (*). **C.** Samples from the experiment shown in A were collected at the indicated time points, proteins were TCA extracted and subjected to western blotting to determine Rad53 phosphorylation during the response to the HO induction. **D.** Cdk-dependent phosphorylation of Psy2 is required for SSA repair but not for PP4-mediated checkpoint inactivation. Western blot of Psy2 to determine its phosphorylation profile in SSA. Cells were grown overnight in YP-Raffinose, and the HO nuclease was induced by adding galactose to the medium. Samples were collected at the indicated time points, and proteins were extracted using TCA precipitation and subjected to Phos-tag western blotting to monitor Psy2 phosphorylation dynamics along the DNA damage response. Coomassie staining is shown as loading control. **E.** Lack of Cdk activity reduces the higher molecular weight isoforms of Psy2 accumulated during the damage response. Cells were HO-expressed for 2 h, DMSO or 1-NM-PP1 was added to the culture and samples were taken two hours from the addition of the drugs. **F.** Illustration depicting the position of Cdk consensus phosphorylation sites within the Psy2 protein sequence. **G.** Southern blot of *rad51Δ* and *rad51Δ psy2-S/T-A* cells carrying the DNA repair system depicted in figure 1A. Cells were grown overnight in YP-Raffinose before inducing the HO nuclease with galactose. Samples were collected at various time points, genomic DNA was extracted, digested with *Kpn*I, and analyzed by Southern blot using the probes indicated in figure 1A. The *ACT1* gene sequence was used as loading control. **H.** Graphs derived from the data obtained in G, representing the quantification of the averaged donor and product band signals from two biological replicates, normalized against their respective *ACT1* and uncut T0 band signals. The intersection point of the donor and product trend lines (I_D/P_ value) represents the mean from two independent experiments. The statistical significance (*P*-value) of differences between both strains assessed by a two-tailed unpaired Student’s *t-*test is 0.0912 (ns). **I.** Samples from the experiment shown in G were collected at the indicated time points. Proteins were TCA extracted and subjected to western blotting to analyze Rad53 phospho-dynamics along the HO induction.

**Expanded View Figure 2.**
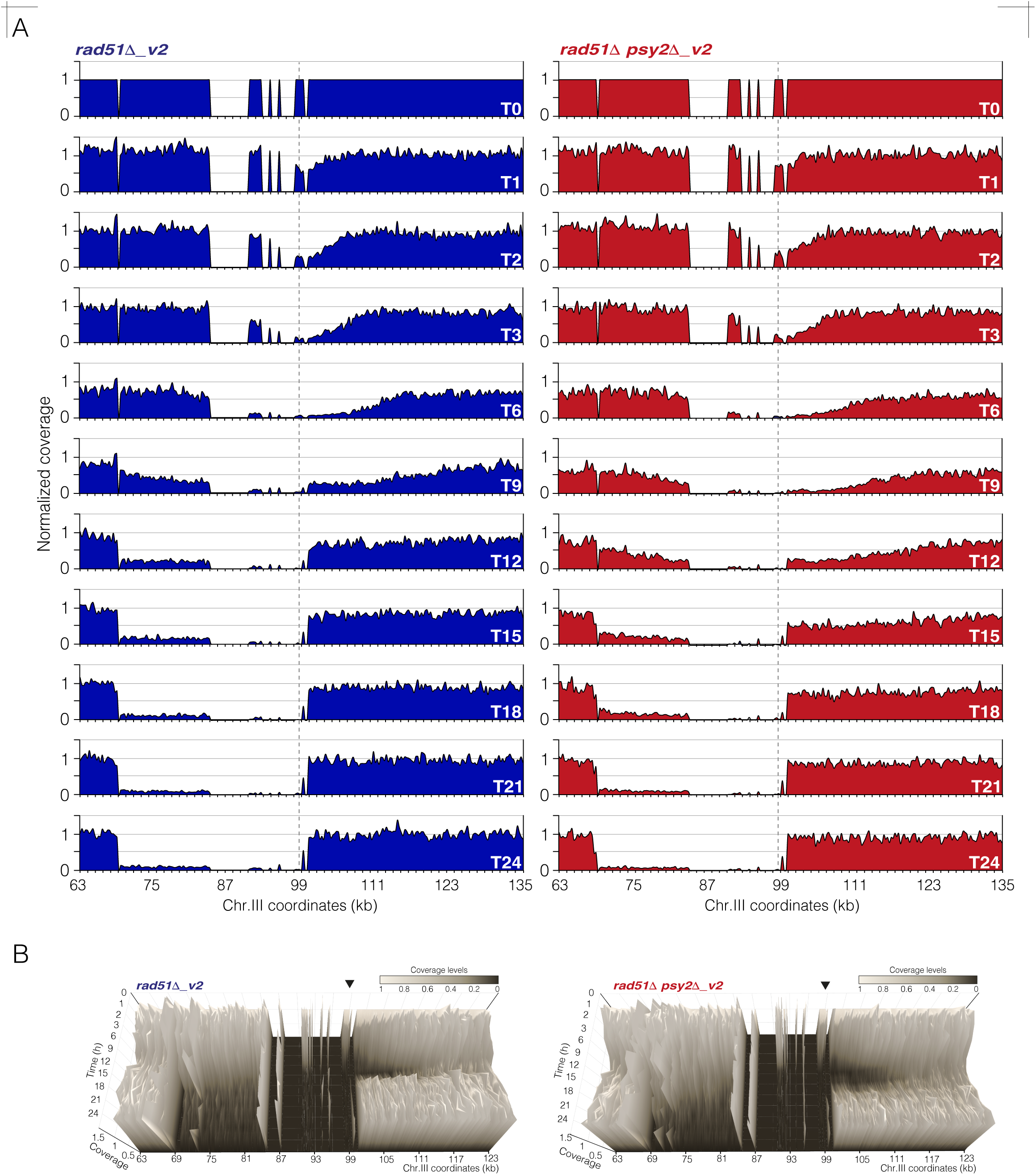
Genome-wide sequencing analysis of SSA repair in *rad51Δ*_v2 and *rad51Δ psy2Δ _v2* strains. **A.** Normalized 2D coverage profiles of *rad51Δ*_v2 and *rad51Δ psy2Δ_v2* cells. Vertical dotted lines mark the position of the HO cleavage sites. The graphs show the means from two biologically independent experiments. **B.** Graphs from A were compiled to generate normalized 3D coverage profiles of *rad51Δ*_v2 and *rad51Δ psy2Δ_v2* cells, simultaneously representing coverage levels (*y-*axis), chromosome III coordinates (*x-*axis), and time after HO induction (*z-*axis). The read coverage values were averaged in 300 nt sections and normalized to their corresponding 0 h sample and to a 120 kb region between coordinates 210000 and 330000 of chromosome V (see Material and Methods for details). White and black represent high- and low-coverage levels, respectively. The black triangles mark the position of the HO break. The graphs show the mean from two biological replicates.

**Expanded View Figure 3.**
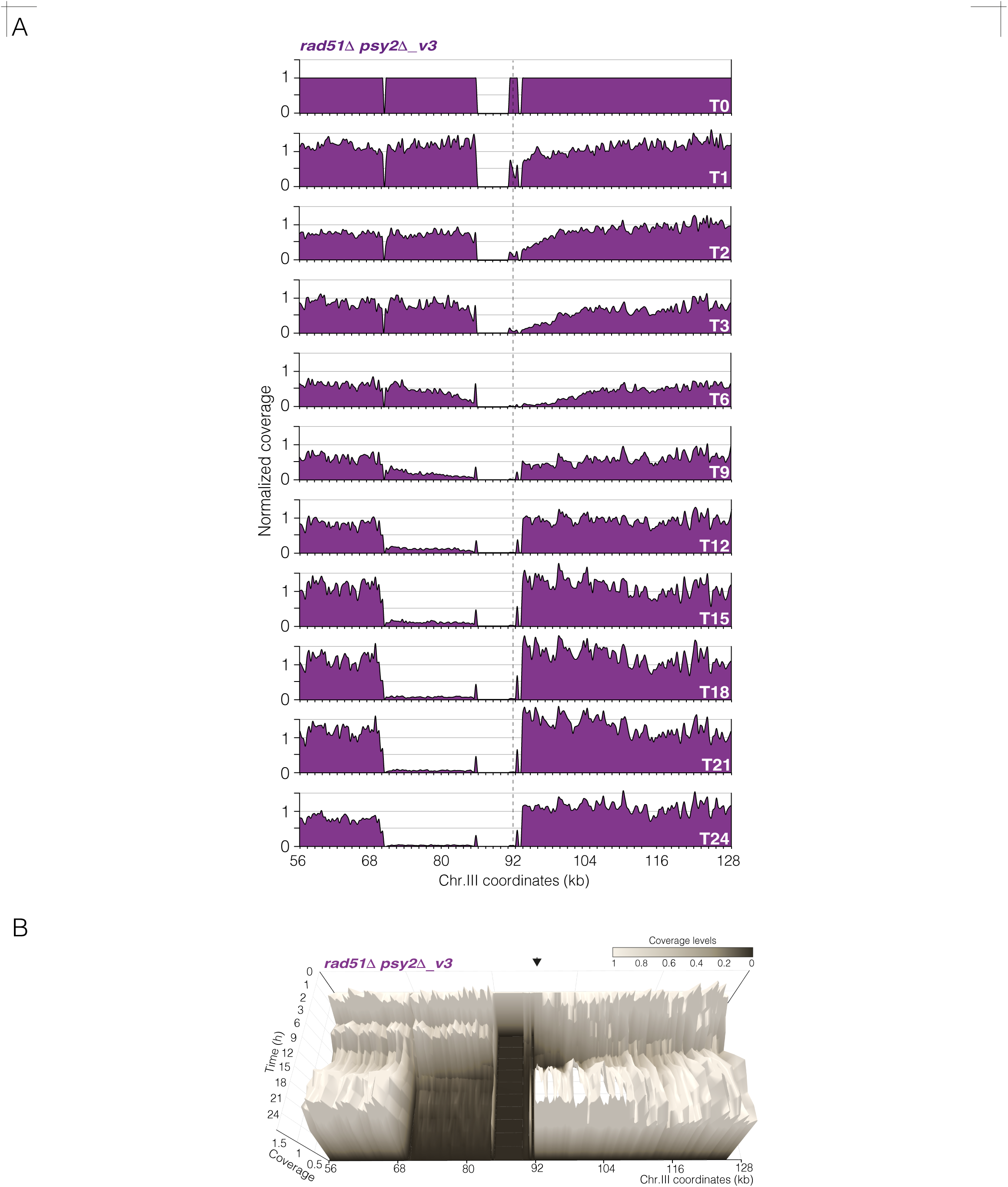
Genome-wide sequencing analysis of SSA repair in the *rad51Δ psy2Δ _v3* strain. **A.** Normalized 2D coverage profile of the *rad51Δ psy2Δ_v3* strain. Vertical dotted line marks the position of the HO cleavage site. The graphs show the means from two biologically independent experiments. **B.** Graphs from A were compiled to generate the normalized 3D coverage profile of the *rad51Δ psy2Δ_v3* strain, simultaneously representing coverage levels (*y-*axis), chromosome III coordinates (*x-*axis), and time after HO induction (*z-*axis). The read coverage values were averaged in 300 nt sections and normalized to their corresponding 0 h sample and to a 120 kb region between coordinates 210000 and 330000 of chromosome V (see Material and Methods for details). White and black represent high- and low-coverage levels, respectively. The black triangle marks the position of the HO break. The graphs show the mean from two biological replicates.

**Expanded View Figure 4.**
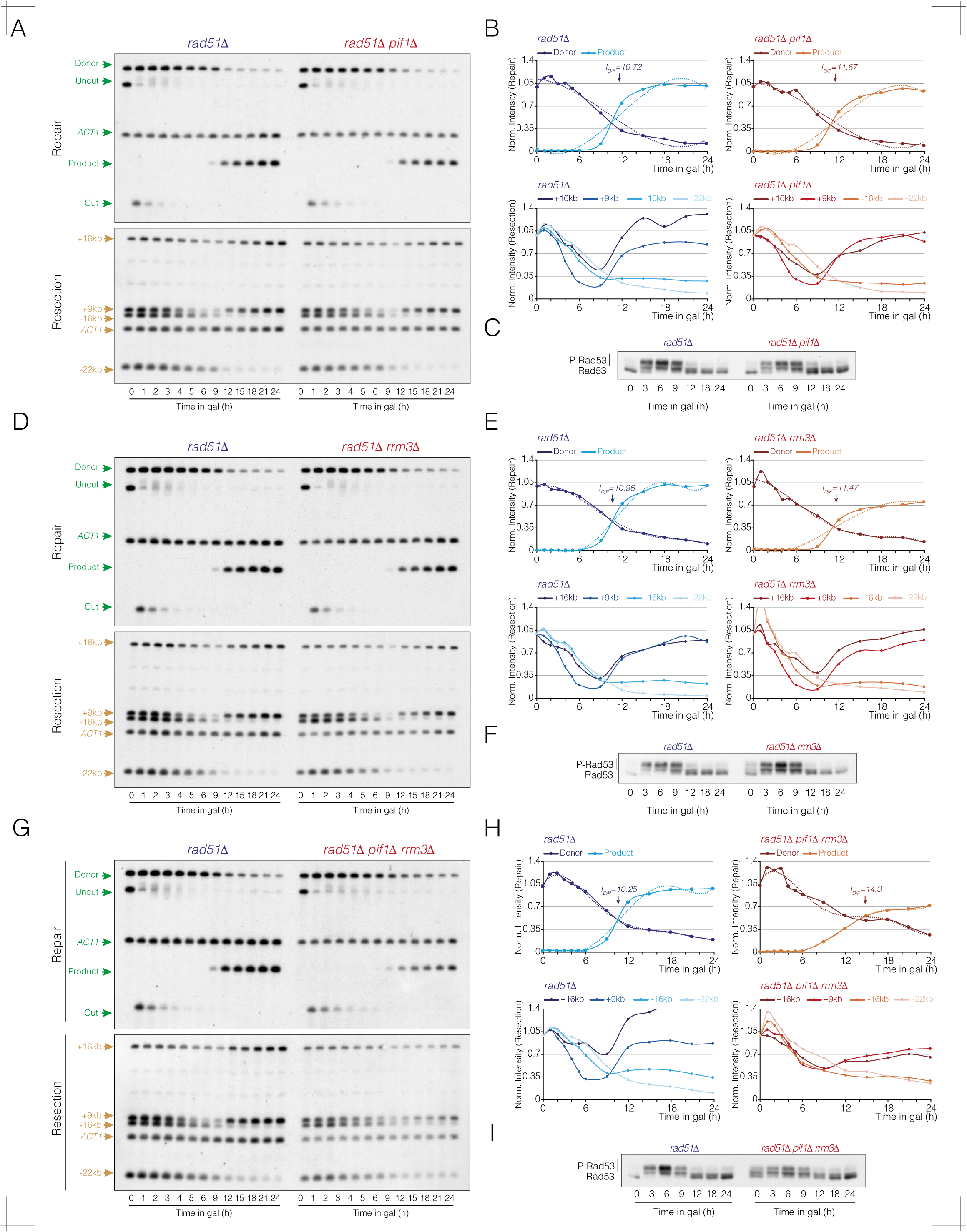
Pif1 and Rrm3 helicases cooperate in SSA by stimulating DNA end resection. **A.** Southern blot analysis of *rad51Δ* and *rad51Δ pif1Δ* strains. Cells were grown overnight in YP-Raffinose before adding galactose to the medium to induce HO expression. Samples were collected at different time intervals, genomic DNA was extracted, digested with *Kpn*I (top panel) or *Sty*I (bottom panel), and subjected to Southern blot analysis using the probes shown in figure 3A. The *ACT1* gene sequence was used as loading control. **B.** Top graphs represent the quantification of the donor and product band signals from A (top panel), averaged from two biological replicates and normalized to their respective *ACT1* and uncut T0 signals. The intersection point of the donor and product trend lines (I_D/P_ value) represents the mean from two independent experiments. The statistical significance (*P*-value) of differences between both strains assessed by a two-tailed unpaired Student’s *t-*test is 0.2587 (ns). Bottom graphs represent the quantification of +16 kb, +9 kb, -16 kb and -22 kb band signal intensities from A (bottom panel), averaged from two independent experiments and normalized to their respective *ACT1* and uncut T0 signals. **C.** Samples from the experiment shown in A were collected at the indicated time points, proteins were TCA extracted and subjected to western blotting to determine the Rad53 phosphorylation profile. **D.** Southern blot analysis of *rad51Δ* and *rad51Δ rrm3Δ* strains using the same experimental conditions as in A. Samples were collected at different time points, genomic DNA was extracted, digested with *Kpn*I (top panel) or *Sty*I (bottom panel), and subjected to Southern blot analysis using the probes shown in figure 3A. The *ACT1* gene sequence was used as loading control. **E.** Top graphs represent the quantification of the donor and product band signals from D (top panel), averaged from two biological replicates and normalized to their respective *ACT1* and uncut T0 signals. The intersection point of the donor and product trend lines (I_D/P_ value) represents the mean from two independent experiments. The statistical significance (*P*-value) of differences between both strains assessed by a two-tailed unpaired Student’s *t-*test is 0.0635 (ns). Bottom graphs represent the quantification of +16 kb, +9 kb, -16 kb and -22 kb band signal intensities from D (bottom panel), averaged from two independent experiments and normalized to their respective *ACT1* and uncut T0 signals. **F.** Samples from the experiment shown in D were collected at the indicated time points, proteins were TCA extracted and subjected to western blotting to determine the Rad53 phosphorylation profile. **G.** Southern blot analysis of *rad51Δ* and *rad51Δ pif1Δ rrm3Δ* strains following the same experimental conditions as in A. Samples were collected at different time points, genomic DNA was extracted, digested with *Kpn*I (top panel) or *Sty*I (bottom panel), and subjected to Southern blot analysis using the probes shown in figure 3A. The *ACT1* gene sequence was used as loading control. **H.** Top graphs represent the quantification of the donor and product band signals from G (top panel), averaged from two biological replicates and normalized to their respective *ACT1* and uncut T0 signals. The intersection point of the donor and product trend lines (I_D/P_ value) represents the mean from two independent experiments. The statistical significance (*P*-value) of differences between both strains assessed by a two-tailed unpaired Student’s *t-*test is 0.0033 (**). Bottom graphs represent the quantification of +16 kb, +9 kb, -16 kb and -22 kb band signal intensities from G (bottom panel), averaged from two independent experiments and normalized to their respective *ACT1* and uncut T0 signals. **I.** Samples from the experiment shown in G were collected at the indicated time points, proteins were TCA extracted and subjected to western blotting to assess Rad53 phosphorylation along the HO induction.

